# Core Microbiota Drive the Prevalence of Extracellular Antibiotic Resistome in the Water Compartments

**DOI:** 10.1101/2022.01.04.474879

**Authors:** Kai-Feng Yu, Peng Li, Yuansheng Huang, Jun Yang, Han Li, Yu Xie, Rui Li, Bo Zhang, Yiliang He

## Abstract

Unlike intracellular chromosome, extracellular DNA (eDNA) may accelerate the spreading of antibiotic resistance genes (ARGs) through natural transformation, but one of the core issues regarding to the bacterial taxonomic characterization of eDNA in the complex water environments is largely unknown. Hence, Illumina Miseq sequencing was used to identify the genotype of eDNA from wastewater (WW), river water (RW) and stormwater (SW) runoff. High-throughput qPCR targeting 384 genes was implemented to detect extracellular ARGs (eARGs) and mobile genetic elements (eMGEs). We obtained 2,708,291 high quality sequences from 66 eDNA samples. The SW exhibited the significant higher Shannon Index. Subsequently, we identified 34 core bacteria sources of eDNA widely distributed in the three water compartments. Among which, *Pseudomonas, Flavobacterium, Limnohabitans, Burkholderiaceae_unclassified, Methylotenera* and *Acinetobacter* were the most prevalent. A total of 302 eARGs and eMGEs were detected, suggesting that eDNA is an important antibiotic resistance reservoir. Among the 127 shared genes of the three groups, 15 core resistance genes were filtered, including *IS6100, sul1* NEW, *intI1, ISPps1-pseud, aac3-Via, qacH_351* and *ISSm2-Xanthob*. The Procrustes analysis and Variance Partitioning Analysis (VPA) demonstrated that core bacteria and MGEs were significantly correlated with eARGs. These results suggested that the occurrence and changes of eARGs in the water compartments may be largely attributed to the core microbiota and eMGEs.

**Synopsis:** The identification of bacterial taxa of extracellular DNA is the preliminary issue for the natural transformation of extracellular antibiotic resistome.

## 1. Introduction

As the counterpart of intracellular DNA (iDNA) located within the cell membranes, extracellular DNA (eDNA) is outside thereof ^1^. eDNA not only originates from spontaneous cell death, autolysis and self-secretion ^2^, but also from the viral infections ^3^, resulting its ubiquitous in the environments. The amount of eDNA was estimated to gigatons globally ^4^ while that in activated sludge was as high as 52 mg per g volatile suspended solids ^5^. It has been demonstrated that eDNA can be long durable in the environment for months and years ^2^. Notably, eDNA can be directly taken up by both Gram-positive and -negative species as carbon and phosphorous sources or for nucleic acid repairing ^6-8^. This natural transformation has been widely accepted as one of the most important mechanisms of horizontal gene transfer (HGT) among microorganisms ^7, 9^. One of the most popular research interests of eDNA have long been located on its role in biofilm formation ^5, 10^. Consequently, eDNA fraction plays a critical role in gene transfer, microbial growth and metabolism activities. However, its role in the spreading of antibiotic resistance genes (ARGs) has not well been recognized.

The possible health risks of ARGs have raised widespread concerns ^11, 12^. Their prevalence in the environments has been broadly uncovered. It has been discovered that intracellular ARGs (iARGs) and intracellular mobile genetic elements (iMGEs) are broadly existed in the diverse environmental compartments, such as wastewater treatment plants (WWTPs) ^13^, river water ^14^, lakes ^15^, soils ^16^ and stormwater runoff ^17^.

The contents of iARGs have been fully enriched, but that of eARGs, especially considering eDNA as carriers of ARGs and MGEs, have only been framed an outline, and remain to be fully understood. It has been revealed that the absolute abundance of eARGs was as high as 3.79 × 10^7^ copies/mL in wastewater ^18^, indicating that wastewater is important eARG reservoir. More importantly, eARGs can be horizontally transferred through natural transformation, which may stimulate the spreading of ARGs ^9^. Efforts have been made to unravel the role of plasmids in the transfer of ARGs through bacterial conjugation. But that of the eDNA fragments is still scarce. To reveal the dynamics of eARGs carried by eDNA in the environment, it is preliminarily urgent to figure out the eDNA fragments from exact which species. At the lab scale, previous studies have demonstrated that strains, such as *Acinetobacter, Bacillus, Flavobacterium, Neisseria* and *Pseudomonas*, can actively secrete DNA ^8, 19^. However, the taxonomic characteristics of eDNA in more complex environments have not been fully uncovered. Most of the bacteria in the wild cannot be cultured at present lab conditions ^20^. Therefore, using high-throughput sequencing technology may help us to get wider pictures of eDNA from natural environments.

Among the diversified environmental compartments, wastewater (WW) has been accepted as a hotspot of ARGs. Riverine water (RW) significantly contributes to the migration and dissemination of ARGs owning to its frequent mobility and connection with other environmental compartments. Stormwater (SW) runoff also driven the dynamics of ARGs ^17, 21^, but the information regarding to eARGs is scarce. Therefore, we selected these three typical water matrices. We hypothesized that there may be shared bacteria between the groups that drive the prevalence of the antibiotic resistome. To profile the bacterial sources of eDNA, and its potential relationship with eARGs, we aimed to 1) identify the core taxa sources of eDNA from the three water compartments, 2) profile the occurrence of eARGs broadly in the WW, RW and SW, and 3) analyze the correlations between the core taxa and eARGs.

## 2. Materials and Methods

### 2.1 Sites description and sampling

A total of 42 sampling sites targeting WW, RW and SW was set (**Table S1**; **Fig. S1**). The WW samples were collected from six full-scale WWTPs in China (See details in **Table S2, Figs. S2** and **S3**, and our previous studies ^22, 23^). One plant (WWTP1) implements a tridimensional eco-biological process with numerous macrophytes planted on the top of the biological reactors. The treated wastewater of the WWTP1 is discharged to a nearby stream. The RW was sampled from the receiving stream (**Fig. S2**). To make the RW samples more representable, we also collected samples from the Yangtze River Basin in China (**Table S1; Fig. S1**). The SW samples were collected from the agricultural and residential zones in Southern China (**Fig. S4**). In addition, considering the low flow rate of the stream in dry spell, SW samples were also obtained from the stream during the stormwater events. The specific sampling strategies are shown as **Text S1** in the Supplementary Information.

### 2.2 Sample pretreatment and eDNA extraction

All of the water samples were initially filtered by a vacuum filter with 0.2-μm pore-size polycarbonate membranes (Millipore, USA). The filters were used to extract iDNA by FastDNA™ Spin Kit for Soil (MP Biomedicals, USA). The filtrates were used for eDNA extractions via ethanol precipitation method ^24-26^. Thirty-three milliliter (mL) 3M sodium acetate solution (Aladdin, Shanghai, China) and 726 mL absolute ethanol (≥ 99.7%; Aladdin) were added into per filtrate sample (330 mL). The mixed solution was stored at -20°C over one night. After that, the mixture was centrifuged for 10 min at 4°C under the speed of 10, 000 × g. The supernatant was discarded, and the pellet precipitate (**Fig. S5**) was air dried and used for DNA extraction by FastDNA™ Spin Kit for Soil (MP Biomedicals, USA) according to the standard protocol.

### 2.3 HT-qPCR

A total of 384 pair primers including major class ARGs and MGEs were designed and listed in **Table S3** according to previous studies ^13, 27^. High throughput qPCR (HT-qPCR) was implemented on SmartChip PCR system including a Multisample Nanodispenser and a Cycler (Takara SmartChip™, previously WaferGen Biosystems). DNA samples were first dispensed, then the mixtures of primers and fluorescence (in 384-well plates) using the Multisample Nanodispenser. The cycling conditions were set according to previous studies ^22, 28, 29^ and shown as follows: 10 min denaturation at 95°C, 40 cycles with 30 s amplification at 95°C and 30 s final extension at 60°C. The amplification efficiencies within 1.8 to 2.2 were considered as positive results. Threshold of 31 cycles (C_t_) was set to be detection limit ^29^. Three replicates were conducted for each sample. Copies of detected ARGs and MGEs were calculated according to the following equation ^30, 31^: 10^(31-Ct)/(10/3)^. Relative abundance of each ARG and MGE was obtained by dividing the copy with 16S rRNA copy. Furthermore, absolute abundance was calculated by multiplying the relative abundance with absolute 16S rRNA copy ^32^ which was measured by real-time quantitative PCR (RT-qPCR) according to plasmid standard curve (**Text S2**).

### 2.4 16S rRNA sequencing

The bacteria 16S rRNA sequencing was conducted by thermocycler PCR system (GeneAmp 9700, ABI, USA). Primers of 338F (5’-ACTCCTACGGGAGGCAGCAG-3’) and 806R (5’-GGACTACHVGGGTWTCTAAT-3’) were used to target V3 - V4 regions. The PCR products were recovered using a 2% agarose gel, purified using AxyPrep DNA Gel Extraction Kit (Axygen Biosciences, Union City, CA, USA) and eluted with Tris-HCl. Detection and quantification using QuantiFluor ™ -ST (Promega, USA). The raw eDNA samples and PCR products were checked by agarose gel electrophoresis (**Fig. S6**). Paired-end sequencing (2 × 300 bp) of purified amplicons was realized on an Illumina MiSeq platform (Illumina, San Diego, USA) based on the standard protocols (Majorbio Bio-Pharm Technology Co. Ltd., China). Quality filtering of raw sequencing data (fastq files) was implemented using Trimmomatic ^33^, and merging by FLASH ^34^. UPARSE software (version 7.1, http://drive5.com/uparse/) was used to perform operational taxonomic unit (OTU) clustering on sequences at the threshold of 97% similarity ^35^. RDP classifier (http://rdp.cme.msu.edu/) was used to annotate the taxonomic classification of each sequence against the Silva database (SSU132) with the alignment threshold of 80% ^28, 36^. The raw sequence datasets were uploaded to the National Center for Biotechnology Information (NCBI) Sequence Read Archive (SRA) with bio-project accessions No. PRJNA592276, PRJNA639397, PRJNA648808, PRJNA687102 and PRJNA729178.

### 2.5 Definitions of core taxa and antibiotic resistome

The rules to filter core taxa of microbiota and antibiotic resistome are according to the previous studies ^37, 38^ and shown as follows: 1) detected in more than 75% samples from all of the three groups; 2) for core taxa alone, there is at least one sample that the relative abundances of one taxon is greater than 1.0%.

### 2.6 Statistical analysis

Statistical analyses of the sequencing data were completed on Majorbio Cloud Platform (https://cloud.majorbio.com/) (Shanghai Majorbio Bio-pharm Technology Co., Ltd, China). Alpha diversity was evaluated by Chaos and Shannon indexes. Beta diversity analysis was by principle coordinate analysis (PCoA; Bray-Curtis distance) followed by Adonis test. When calculating the relative abundances of taxa and conducting PCoA, all the sample sequences were normalized to the least sequence read depth ^39^. Multiple group diversities of the bacterial communities were checked using Kruskal-Wallis (K-W) test followed by Tukey-kramer post-hoc test. P values were corrected by false discovery rate (FDR) method.

The analysis of variance (ANOVA, one-way, turkey post-hoc test) was conducted using SPSS 21.0 (IBM, USA). Principle coordinate analysis of eARGs and redundancy analysis (RDA) based on Bray-Curtis distance were implemented by using Canoco V5.0 software ^40^. One-way permutational multivariate analysis of variance (PERMANOVA, Bray-Curtis distance, 9999 permutations) was performed by PAST V4.0.1 ^41^. Statistic *p*-value less than 0.05 was considered significant. Heatmap was visualized by HemI 1.0. Veen diagrams were plotted by VENNY 2.1 (https://bioinfogp.cnb.csic.es/tools/venny/index.html). Origin 2018 (Origin Lab, USA) was used to draw the char and box charts. Network analysis was conducted by Gephi 0.9.2.

## 3 Results

### 3.1 Taxonomic Characterization of eDNA

#### 3.1.1 Structures of microbial communities

A total of 2,708,291 high quality sequences were obtained from 66 eDNA samples (**Table S4**). The sequence numbers of eDNA were comparable with iDNA (**Table S5**). Subsequently, 4,906 OTUs were obtained. Among which, 627 OTUs were shared while 3,174, 168 and 132 OTUs were unique among the WW, RW and SW, respectively (**Fig. 1a**). The WW had the most OTU number (4521) compared to the other two compartments. But the SW had the highest Shannon Index (**Fig. 1b; Table S4**). These results indicated that the eDNA source microorganisms were rich in diversity, especially in the SW and WW. The eDNA belonged to the classes of *Gammaproteobacteria, Bacteroidia, Alphaproteobacteria* and *Verrucomicrobiae* were the most abundant in the three groups (**Fig. 1c; Fig. S7**). At genus level, the PCoA result (Bray-Curtis distance; **Fig. 1d**) indicated that the microbial communities of the three groups were significantly different (Adonis test, *r* = 0.1110, *p* = 0.0001).

**Fig. 1.**
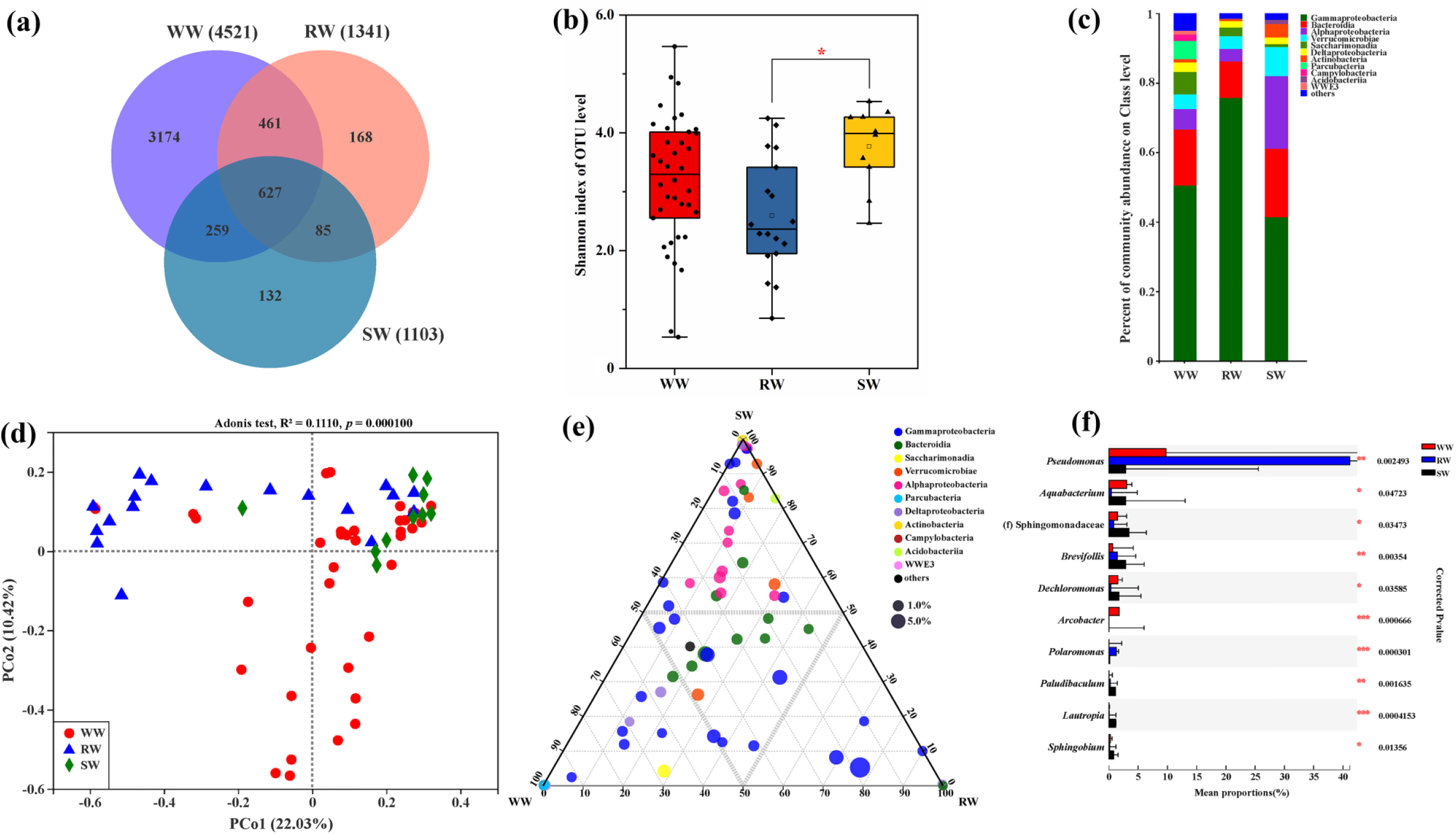
The taxa characters of eDNA are presented. The Venn plot (**a**) displays the shared and unique OTU numbers in the wastewater (WW), riverine water (RW) and stormwater runoff (SW). The box plots present the Shannon index of the WW, RW and SW (**b**). The relative abundances of microbial communities of the WW, RW and SW on Class level (**c**). Bray-Curtis distance-based principle coordinates analysis (PCoA) reveals the distribution patterns of the genera (relative abundances) from the three water compartments (**d**). The ternary diagram shows the dominant and distribution of genera among the WW, RW and SW (**e**). The symbols present the genera, and colors are ranked according to the corresponding classes. Circle size means average abundance. Comparisons of bacterial differences (genus level) between the three groups were conducted by Kruskal-Wallis (K-W) test followed by Tukey-kramer post-hoc test (**f**; *, *p* < 0.05; **, *p* < 0.01; ***, *p* < 0.01).

The tertiary plot showed the distribution patterns of genera among the WW, SW and RW (**Fig. 1e**). The bacteria in the middle (thick dash line) of the tertiary plot were commonly shared between groups, such as *Flavobacterium, Limnohabitans, Burkholderiaceae_unclassified* and *Acinetobacter*. The details were presented in **Table S6**. The rare genera were close to the vertexes of the triangle. Furthermore, the multiple group comparison at genus level was performed to reveal the differences of microbial communities (**Fig. 1f**). Among the top 10 significantly different genera (K-W tests, FDR corrected *p* < 0.05), *Pseudomonas* and *Polaromonas* in the WW exhibited the higher relative abundances than that in the others while *Aquabacterium* and *Dechloromonas* were more prevalent in both the WW and SW. But *Arcobacter* was abundant alone in the WW. The rest genera were more prevalent in the SW.

#### 3.1.2 eDNA from core microbiota

Among the prevalent bacteria (genus level; **Fig. S8**), we also identified 34 core bacteria from the 66 eDNA samples (**Fig. 2a**). Extracellular DNA from the genera of *Pseudomonas* (17.25%), *Flavobacterium* (6.32%), *Limnohabitans* (5.21%), *Burkholderiaceae_unclassified* (5.17%), *Methylotenera* (4.84%), *Acinetobacter* (4.43%), *Saccharimonadales_norank* (4.10%) and *Aquabacterium* (2.27%) exhibited higher prevalence. It should be noted that *Acinetobacter* and *Burkholderiaceae_unclassified* were 100% detected. These results suggested that the core bacteria were potentially the most commonly found sources of eDNA in the water matrixes. To further confirm these results, we also profiled the taxa of the corresponding iDNA samples (**Table S5; Fig. S9**). All of the core bacteria were detected in the iDNA samples while 26 out of 34 strains were also classified to core (**Fig. S10**), suggesting that eDNA were from the prevalent source bacteria.

**Fig. 2.**
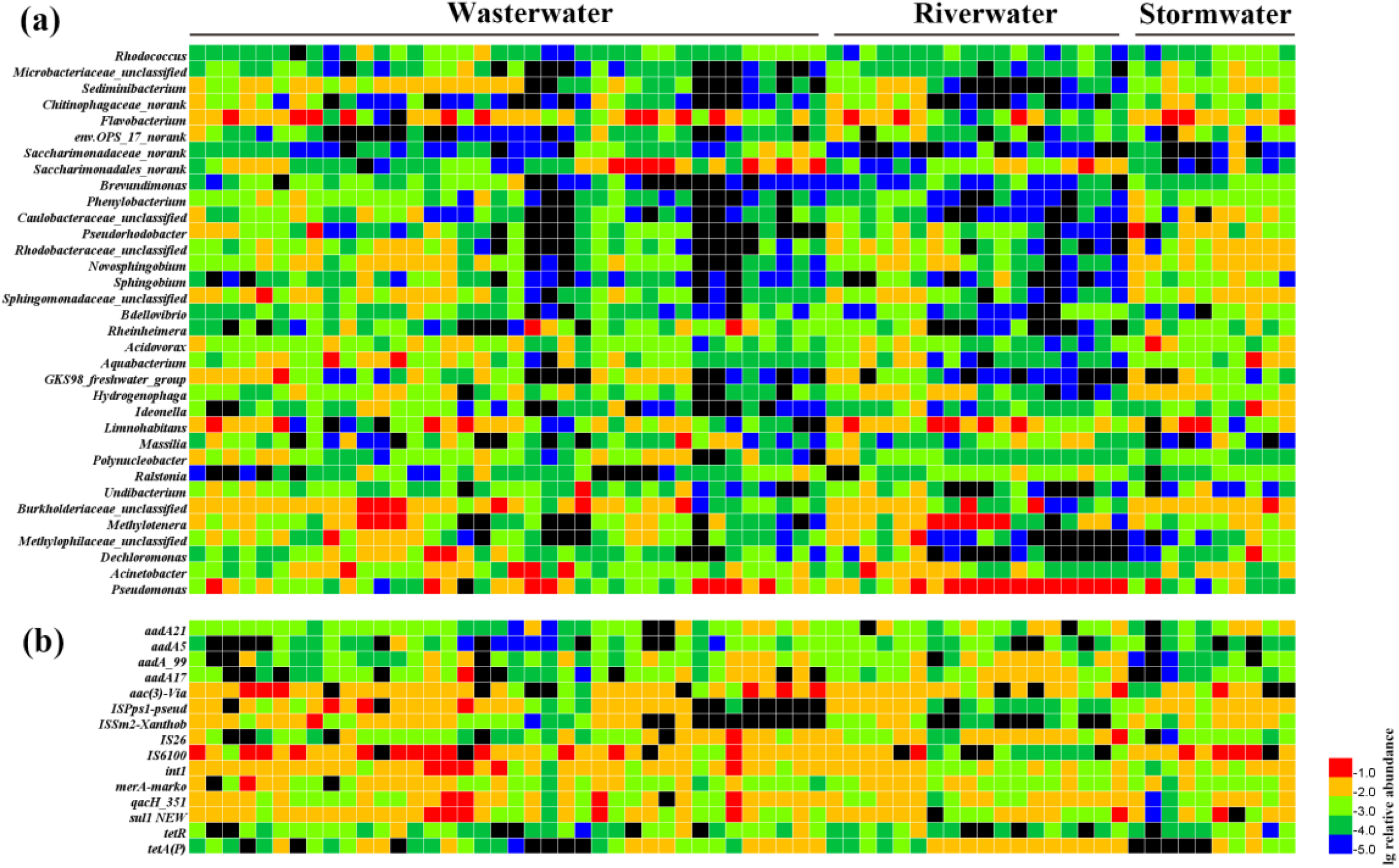
The heatmaps show the relative abundances (lg transformed) of core bacteria (genus level; **a**) and antibiotic resistome (**b**) from the eDNA samples in the WW, RW and SW.

### 3.2 Diversity and Occurrence of extracellular ARGs and MGEs

#### 3.2.1 Number of eARGs and eMGEs

On average, 96 out of 383 ARG and MGE types were detected in the RW (**Fig. 3a**), which is significantly higher than that of the WW and SW (one-way ANOVA, *p* < 0.01). The lowest number of eARGs and eMGEs were obtained in the SW. Of the 13 major resistance classes, Aminoglycoside, MGEs, multidrug resistance (MDR) and MLSB resistance genes were the most detected types in the three groups. Our results suggested that eARGs and eMGEs universally existed in the water compartments. Among the total detected 302 types of ARGs and MGEs, 127 genes were shared (**Fig. 3b**), while 55, 18 and 4 genes were unique in the WW, RW and SW, respectively. The exact shared and unique genes were illustrated in **Fig. S11**. Cluster I with 127 shared genes among the three compartments should be of the most concern.

**Fig. 3.**
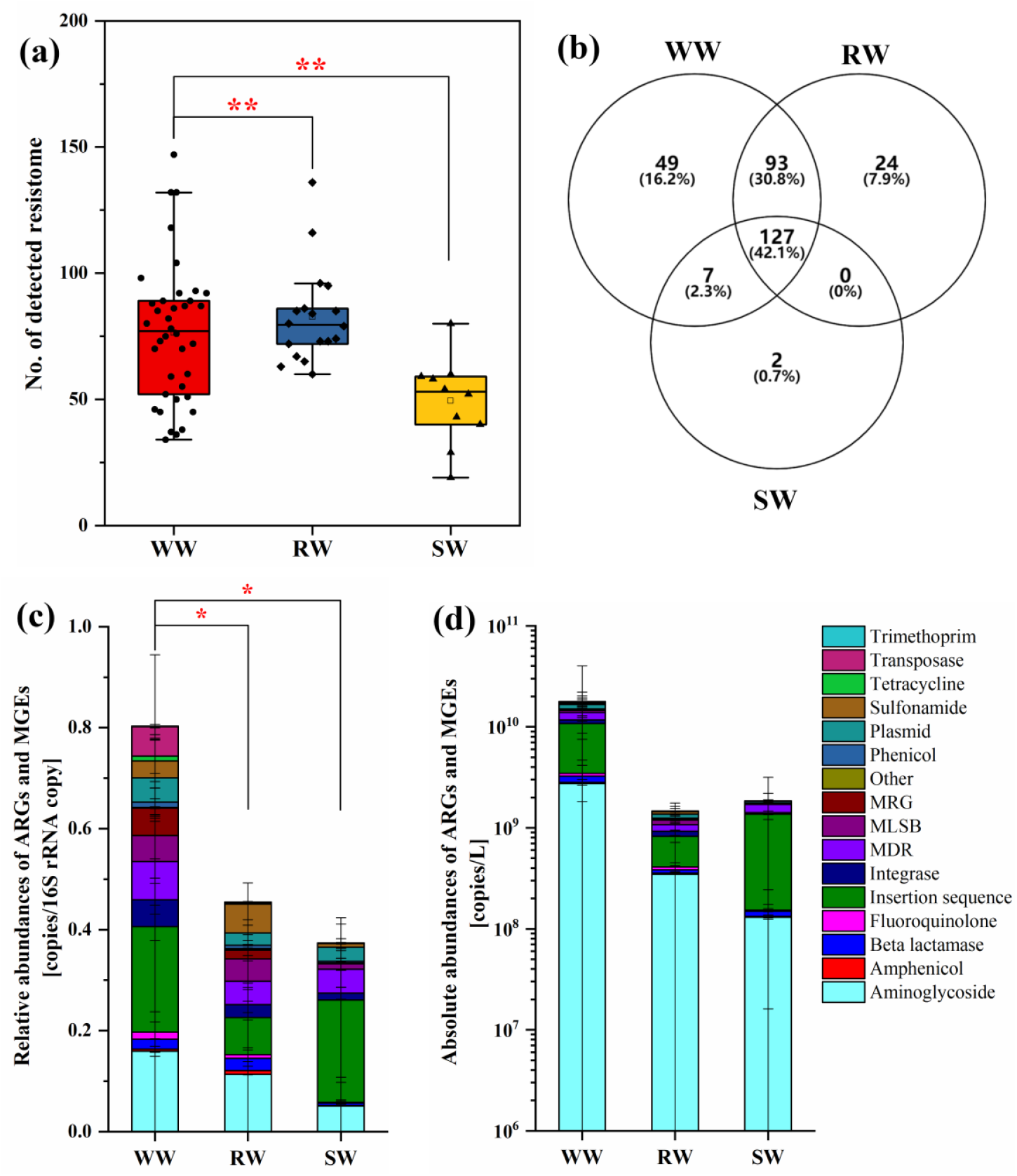
Number of detected eARG and eMGE subtypes in the WW, SW and RW (**a**). The Venn diagram showed the shared and unique numbers of eARGs and eMGEs in the WW, SW and RW (**b**). The bar plots display the relative (**c**) and absolute (**d**) abundances of eARGs and eMGEs. Their subtypes are color ranked. The significant differences are marked (*, *p* < 0.05; **, *p* < 0.01)

#### 3.2.2. Abundance of eARGs and eMGEs

The average relative abundances of eARGs were 0.95, 0.62 and 0.53 in the WW, RW and SW (**Fig. 3c**), respectively. That of eMGEs were 0.35, 0.20 and 0.25(**Fig. 3c**), respectively. The total relative abundance of eARG and eMGE of the WW was significantly higher than that of the RW and SW (one-way ANOVA, *p* < 0.05), respectively. These results further demonstrated that the WWTPs are hotspots of ARGs.

The relative abundance of each gene was presented in **Fig. S12**. The result of PCoA with two axes contributing 36.26% variances showed that the relative abundances of extracellular resistome were significantly different among the three kinds of water (**Fig. 4**; one-way PERMANOVA, *F* = 4.81, *p* = 0.0217), indicating that the characterization of the compartment has the potential to influence the structural dynamics of eARGs and eMGEs.

**Fig. 4.**
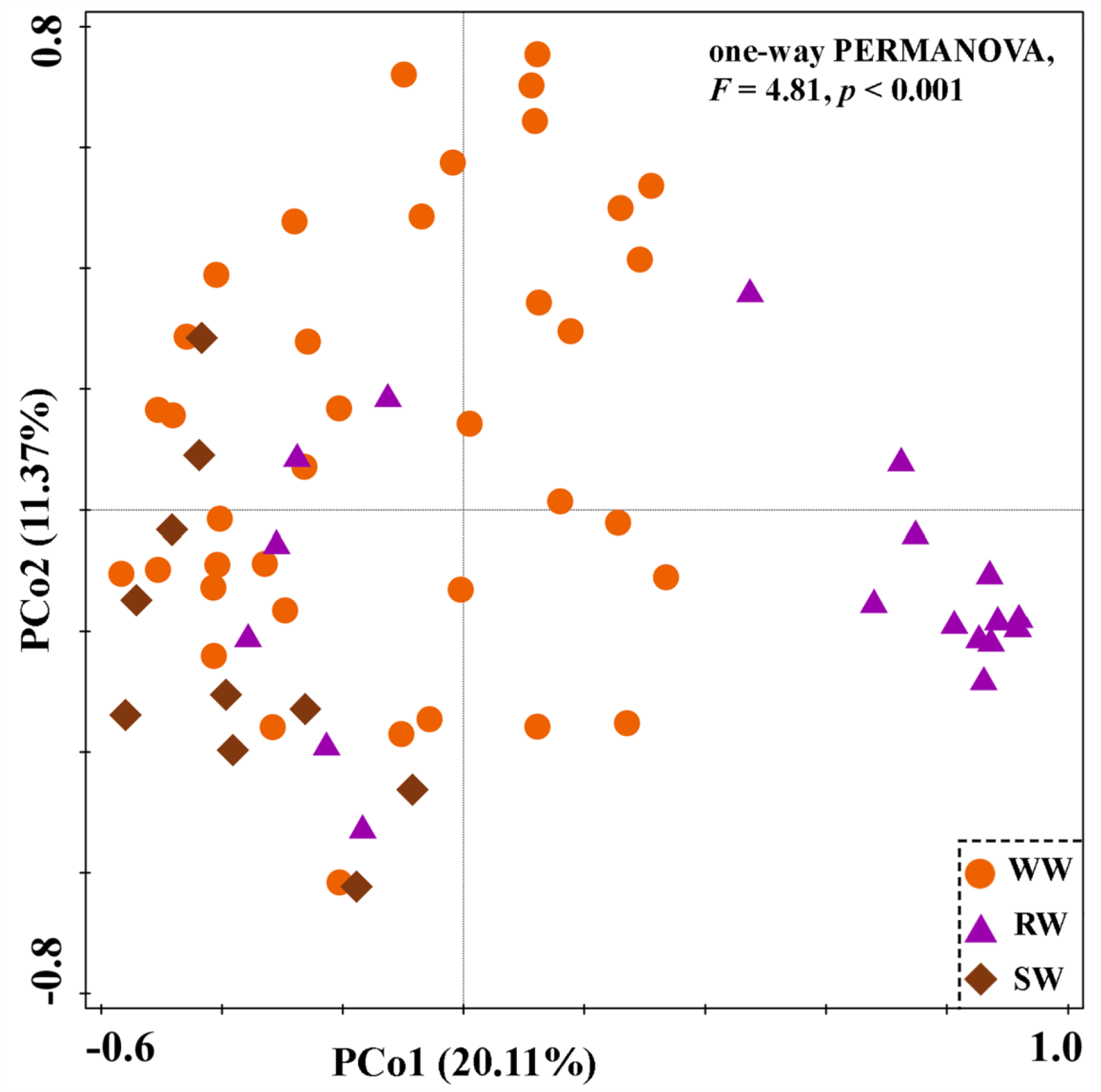
Principal coordinate analysis (PCoA) based on the Bray-Curtis distance showed the overall distribution patterns of eARGs and eMGEs in the WW, RW and SW.

The WW had the most abundant eARGs and eMGEs (**Fig. 3d**), suggesting that the WWTPs are the most important reservoirs of eARGs and eMGEs. We can possibly classify the WWTPs as the hotspots of eARGs. The total absolute abundances of eARGs and eMGEs in the RW and SW were (1.46 ± 2.54) ×10^9^ copies/L and (1.85 ± 2.27) ×10^9^ copies/L, respectively (**Fig. 3d**). No significant differences were observed between the three groups (K-W test, *p* > 0.05). These results revealed that eARGs and eMGEs were prevalent in all the three water compartments.

#### 3.2.3 Core antibiotic resistome

We identified 15 core resistance genes from the 66 eDNA samples (**Fig. 2b**). In detail, there were 5 genes conferring resistance to Aminoglycoside (*aad*A21, *aad*A5, *aad*A_99, *aad*A17 and *aac*(3)-Via), 4 insertional sequences (*IS26, IS6100, ISPps1-pseud* and *ISSm2-Xanthob*), 2 MDR (*merA-marko* and *qacH_351*), 2 Tetracycline (*tetR* and *tet(A)P*), 1 integron (*intI1*) and 1 gene conferring resistance to Sulfonamide (*sul1* NEW). The genes of *IS6100, sul1* NEW, *intI1, ISPps1-pseud, aac3-Via, qacH_351* and *ISSm2-Xanthob* were the most abundant. Our results support an important fact that core eARGs and eMGEs are widely distributed in the water compartments.

### 3.3 Relationship between microbial communities and eARGs

Procrustes analysis demonstrated that the taxa of eDNA was significantly correlated with eARGs (sum of squares M^2^ = 0.5337, *p* < 0.001, 9999 permutations; **Fig. 5a**). Mantel test further confirmed the significant correlations (r = 0.5006, *p* < 0.001). Bray-Curtis distance-based RDA was conducted to reveal the correlations between eARGs and eDNA of source bacteria at genus level (**Fig. 5b**). Thirteen core genera (arrows) exhibited significantly correlations with eARGs and eMGEs (pseudo-*F* = 2.5, *p* < 0.001). Among which, *Saccharimonadales_norank* and *GKS98_freshwater_group* were significantly correlated with eARGs and eMGEs in the WW. In the RW, eARGs were more closely correlated with *Pseudomonas* and *Methylotenera*. The rest of the genera exhibited significantly correlations with eARGs and eMGEs in the WW and SW. These statistical correlations suggested that these core bacteria exhibited great potential to contribute the diversities of eARGs and eMGEs. Furthermore, the VPA results demonstrated that the 23.2% variances of the changes of eARGs were explained by the core bacteria (**Fig. 5c**). In addition, the network analysis indicated that 6 core genera were significantly correlated with resistance genes (**Fig. 6**). A total of 27 MGEs were in the 154 nodes. These results further suggested that the occurrence of eARGs can be attributed to the core bacteria and MGEs.

**Fig. 5.**
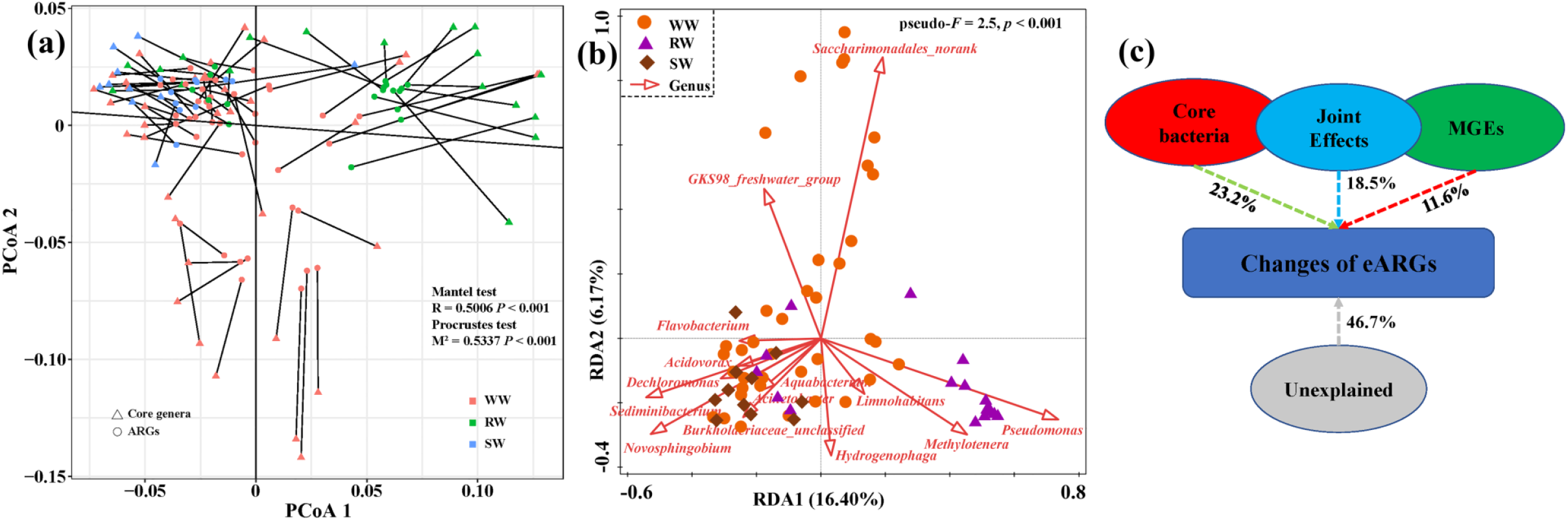
Procrustes analysis shows the significant correlation between antibiotic resistance genes and taxa of eDNA (genus level) based on Bray-Custis distance (sum of squares M^2^ = 0.5337, *p* < 0.001, 9999 permutations; **a**). Mantel test also demonstrates the significant correlation (R = 0.5006, *p* < 0.001). Bray-Custis distance-based redundancy analysis (db-RDA) illustrated the correlation of ARG and MGE subtypes with bacterial community (genus level) (**b**). Thirteen bacteria strains were significantly correlated with the distribution of eARGs (*pseudo*-F = 2.5, *p* < 0.001). Variance partitioning analysis (VPA) was implemented to reveal the contribution of core genera and MGEs to the changes of eARGs (**c**).

**Fig. 6.**
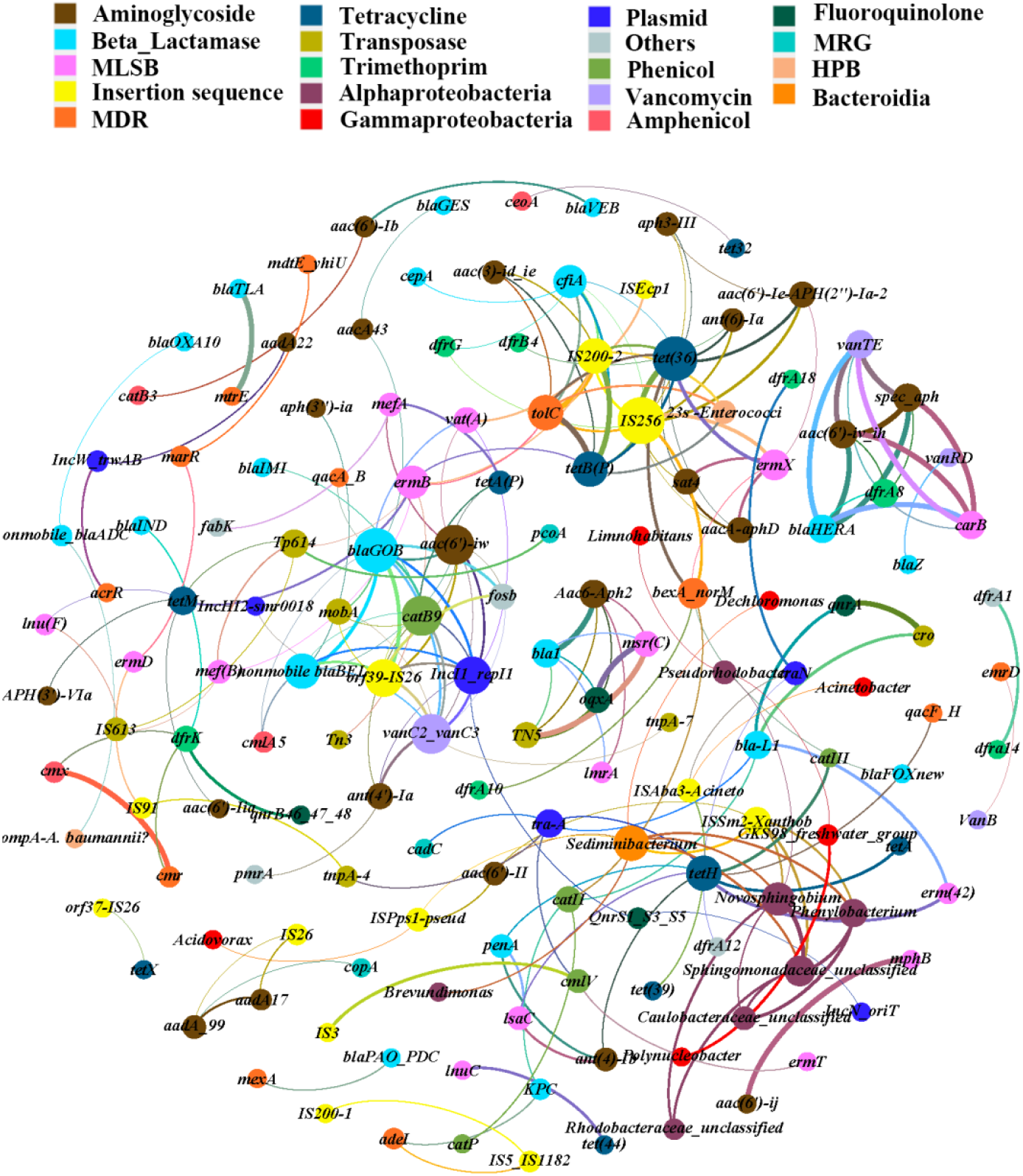
Cooccurrence patterns of eARGs, eMGEs and core genera from all of the eDNA samples were illustrated. A connection represents a significant correlation (Spearman’s co-efficiency ≥ 0.65, *p* < 0.01). The nodes were colored according to ARG and MGE subtypes, and corresponding class level. The node size is ranked according to the number of connections (degree).

## 4 Discussion

### 4.1 Core microbiota as the potential sources of extracellular antibiotic resistome

The release of eDNA from bacteria is a general rule, but what we concerned is that eDNA from which strains can be prevalent and contribute to the dissemination of ARGs. The analysis of core microbiota may address this issue to some extent. On average, the high-quality sequence numbers of eDNA (**Table S4**) were comparable with iDNA (**Table S5**), suggesting that eDNA is an important gene pool. It should also be noted that not all DNA can be prevalent as a extracellular components. For the class level, *Gammaproteobacteria, Bacterodia, Alphaproteobacteria* and *Verrucomicrobiae* have been demonstrated to be widely distributed in the WW ^42, 43^, SW ^44, 45^ and RW ^31, 39^. Subsequently, *Pseudomonas, Flavobacterium, Limnohabitans, Burkholderiaceae_unclassified, Methylotenera* and *Acinetobacter* within those classes were ubiquitous in the water compartments. These abundant bacteria had the greatest potentials to release the corresponding eDNA (**Fig. 2a**) during the decline phase. Obviously, the more bacteria, the more eDNA. In addition, the prevalent core microbiota of eDNA may be largely attributed to the active secretion of defined genera, such as *Pseudomonas, Flavobacterium* and *Acinetobacter* ^2, 9, 46, 47^. Baral et al. found that *Pseudomonas* was the most prevalent in the SW ^45^. Our results (**Fig. 2a**) were in consistent with the previous findings. Notably, disinfection processes as the last defending line of the WWTPs potentially stimulate the release of eDNA ^48, 49^. Although eDNA can be degraded or hydrolyzed, there have been protection matrix formed by particulate materials or dissolved organic matters (DOM) ^50, 51^, resulting their abundance in the environment. Taking together, there are critical pathways that leads to the diversity and prevalence of eDNA in water matrices.

Previous study also showed that the eDNA fragments remained biological intact during the waste disposal processes ^52^. Owing to this significant trait of eDNA, it is reasonable to profile eDNA using high-through put PCR. It has been also demonstrated that the eDNA size up to 1, 700 bp can be sequenced via PCR ^2^. Therefore, using Illumina MiSeq to obtain the taxonomic characterization of eDNA fragments is reliable. As eDNA fragment sizes were located from 80 to more than 20, 000 bp ^1^, 2 × 300 bp paired-end sequencing (**Fig. S6**) may technologically neglect those small length fragments. However, previous study indicated that eDNA smaller than 200 bp only accounted for a very low proportion of total eDNA ^53^. Therefore, our results covered most of the taxonomic information of eDNA.

Dynamics of microbial communities have been accepted as one of the main contributors for the changes of ARGs in natural environments. Our results (**Fig. 5b**) further confirm this conclusion. In addition, water matrices, especially wastewater, are hotspots of ARB ^54, 55^. Therefore, the release of diverse and prevalent of eARGs and eMGEs from those ARB encoded with genes conferring resistances to commonly used antibiotics were spontaneously occurred. Specifically, the strains of *Acinetobacter* have been identified for their role in the release of eARGs, such as *adeK, mdfA* and *mdtK* ^45^. *Flavobacterium* carried diverse genes conferring resistance to Aminoglycosides, Sulfonamides and Tetracyclines. It has also been discovered that *Burkholderiaceae_unclassified* are important source bacteria of ARGs ^56^. *Pseudomonas* strains has commonly known for their vehicle role in disseminating diverse ARGs ^57^. Therefore, the identification and separation of those core bacteria are important to understand the occurrence and discrimination of eARGs owning to their persistence and transformable in the water matrices, which deserves further profiling beyond the first step of recognizing the taxa traits of eDNA.

### 4.2 The Prevalence of Core eARGs and eMGEs

Among the detected eARGs and eMGEs, the core resistome may be the most concerned. The identified core antibiotic resistome (**Fig. 2b**) can be widely distributed, and eMGEs potentially play an important role in the proliferation of ARGs. As to the fact that the aminoglycoside resistance genes have been detected in intercellular samples from typical environmental compartments ^31^, it is no surprising that these genes were commonly found in the extracellular fractions ^58^. In addition, the core genes resistant to aminoglycoside, such as *aadA2-1, aadA5* and *aadA17*, are generally encoded by plasmids, transposons and integrons, which may facilitate their disseminations through HGT ^59^. The genes of *int*I1 and *sul1* have been considered as indicators for anthropological activities ^60, 61^, which is corresponding to our sampling sites significantly influenced by human beings. It is no surprising that *int*I1 is one of the most prevalent MGEs. In addition, these core resistance genes are prone to be carried by plasmids. The mobile elements (integron, IS and plasmids) have been demonstrated to be conducive to the transduction and transformation of ARGs. Numerous studies have confirmed that bacterial cells can recruit eDNA from the surrounding ^7, 62^. The extracellular gene pool should not be neglected. Considering the distribution and mobility characterizations, we can probably put the core antibiotic resistome shared by the three compartments into the priority control list.

### 4.3 Environmental implication

Taking the persistence and transformable traits of eDNA and the mobility of surface water into account, eDNA has the great potential to stimulate the transfer and proliferation of ARGs. Although great efforts have been done to unravel the natural transformation of eDNA in the lab ^47, 62^, more attentions should be paid to the fields where the situations are much more complicated. As one step of the Long March for tackling antimicrobial resistance (AMR), recognizing the diversity and occurrence of eARGs and eMGEs and core taxa sources in the complex environments is very important for us to understand and control the dissemination of ARGs through extracellular sources, which is generally neglected ^24^. Our results reveal the shared core extracellular resistome and possible sources from the core microbiota in the typical water matrices. But these are not enough for the Long March to control the threats of AMR. In the future, attempts can be made to profile the transformation frequency of eARGs between uncultured species, and develop efficient processes to remove eARGs and eMGEs during wastewater treatment.

## Acknowledgments

We sincerely thank Professor Yong-Guan Zhu and Jian-Qiang Su for their great help to provide us the high throughput qPCR platform for the initial measurements and data analyses. Our work was financially supported by National Key R&D Program of China (2019YFD1100202).

## Supporting Information

Text S1, Sampling strategies for the three water compartments; Text S2, Real-time quantitative PCR; Table S1, Descriptions of sampling sites; Table S2, Descriptions of the six full-scale WWTPs; Table S3, Primer sets of the targeted ARGs and MGEs; Table S4, Sequence number and diversity indexes of the eDNA samples; Table S5, Sequence number and diversity indexes of the iDNA samples; Table S6, Results of the tertiary plot; Figure S1, Sampling sites of the riverine water, wastewater and stormwater runoff; Figure S2, The treatment processes of the six full-scale WWTPs; Figure S3, The aerial views of the six WWTPs; Figure S4, The pictures show the sampling sites of stormwater runoff; Figure S5, The obtained extracellular precipitates; Figure S6, Gel electrophoresis images of eDNA and corresponding PCR products; Figure S7, The relative abundances of microbial communities of all eDNA samples on Class level; Figure S8, The heatmap shows the relative abundances (lg transformed) of the abundant genera of eDNA samples; Figure S9, The relative abundances of microbial communities of the iDNA and eDNA samples on Class level; Figure S10, The heatmaps show the relative abundances (lg transformed) of bacteria (genus level) from the iDNA samples; Figure S11, Bipartite network analysis presents the exact shared and unique resistance genes among the three water compartments; Figure S12, The heatmap shows the relative abundances (lg transformed) of all detected extracellular resistance genes;

