## supplementary information for "Core Microbiota Drive the Prevalence of Extracellular Antibiotic Resistome in the Water Compartments"

Supplementary Information contains **45 pages, 2 texts, 6 tables and 12 figures**.

**Text S1** Sampling strategies for the three water compartments.

**Text S2** Real-time quantitative PCR.

**Table S1** Descriptions of sampling sites.

**Table S2** Descriptions of the six full-scale WWTPs.

**Table S3** Primer sets of the targeted ARGs and MGEs.

**Table S4** Sequence number and diversity indexes of the eDNA samples.

**Table S5** Sequence number and diversity indexes of the iDNA samples.

**Table S6** Results of the tertiary plot.

**Figure S1** Sampling sites of the riverine water, wastewater and stormwater runoff.

**Figure S2** The treatment processes of the six full-scale WWTPs.

**Figure S3** The aerial views of the six WWTPs.

**Figure S4** The pictures show the sampling sites of stormwater runoff.

**Figure S5** The obtained extracellular precipitates.

**Figure S6** Gel electrophoresis images of eDNA and corresponding PCR products.

**Figure S7** The relative abundances of microbial communities of all eDNA samples on Class level.

**Figure S8** The heatmap shows the relative abundances (lg transformed) of the abundant genera of eDNA samples.

**Figure S9** The relative abundances of microbial communities of the iDNA and eDNA samples on Class level.

**Figure S10** The heatmaps show the relative abundances (lg transformed) of bacteria (genus level) from the iDNA samples.

**Figure S11** Bipartite network analysis presents the exact shared and unique resistance genes among the three water compartments.

**Figure S12** The heatmap shows the relative abundances (lg transformed) of all detected extracellular resistance genes.

---

Correspondence: Yiliang He, PhD, School of Environmental Science & Engineering, Shanghai Jiao Tong University, 800 Dongchuan Road, Shanghai 200240, China. Tel.: 86-21-54744008.

**Text S1** Sampling strategies for the three water compartments:

***1. Wastewater***

Wastewater treatment plant (WWTP) has been widely accepted as a hotspot of antibiotic resistance genes (ARGs) [1]. It deserves much more attention. Therefore, we gathered 38 wastewater samples in total. These samples were collected from six full-scale WWTPs (**Figs. S2** and **S3**) in China. The selected WWTPs are distributed both in North and South (**Fig. S1**). Among which, WWTP1 is our resident experimental base. To achieve higher discharge standards (third level of *Environmental Quality Standards for Surface Water*), the tridimensional eco-biological treatment processes with numerous macrophytes planted was applied in the plant. Therefore, we sampled along the treatment processes within 2 years (from Jan. 2019 to Sep. 2020) to evaluate the dynamics of ARGs in different treatment units. WWTP2 was selected as the control plant in the same city of WWTP1. The other four WWTPs were randomly selected. Only two sampling sites, including raw influent and effluent of second sedimentation tank, were set in each of the four WWTPs (**Fig. S2**). The descriptions of the wastewater sampling sites are listed in **Table S1**. The detailed information of the six WWTPs are listed in **Table S2**.

***2. Riverine water***

The riverine water samples were collected from two regions, Gaopu Stream and Yangtze River Basin (**Fig. S1**). The stream is the receiving stream of discharged effluent of WWTP1. The average flow rate of the stream is 0.8 $m^3/s$. We set four sampling sites from upstream to downstream. The water samples were collected both in dry season (Jan. 2019) and wet season (July 2019). To make sampling more representative, we also collected riverine water from the Yangtze River Basin. The sampling sites were set along the flow direction, from the Three Gorges Reservoir located in Yichang City to the estuary in Shanghai City (**Table S1**). Nine sampling sites are in the main stream while 2 sites are located in the two main tributaries (Xiangxi River and Hanjiang River). The rest site is the connect channel of Dongting Lake and Yangtze River. The farthest sampling points are approximately 1000 kilometers apart.

### *3. Stormwater runoff*

The stormwater samples were collected from Southern China. The sampling sites were set according to functional districts, including residential area, agricultural zone and industrial district (**Fig. S1**). Considering the fact that the stream is rain-source type and its average flow is 0.8 $m^3/s$, we also conducted 2 sampling sites in the stream (**Fig. S4**). Stormwater runoff sample was collected according to time line ranking from 0 (runoff beginning), 5 min, 10 min, 15min, 20 min, 30 min, 40 min and 1 h. We monitored two stormwater events in Heyuan City, Guangdong Province. The rainfall intensity of Event 1 was 64 mm per 12 h while that of Event 2 was 43 mm per 12 h.

The collected water samples were transported to lab on ice as soon as possible. To avoid abnormal excretion of extracellular DNA (eDNA) during sampling, the water samples were filtered through 0.2-μm pore-sized polycarbonate membranes (Millipore, USA) within 8 hours. The obtained filtrates were used to extract extracellular DNA.

**Text S2** Real-time quantitative PCR

A LightCycler 480 System (Roche) was applied to RT-qPCR to quantify the 16S rRNA according to the plasmid standard curve. Each sample was analyzed in triplicate with a reaction volume of 20 μL, including 10 μL 2×LightCycle 480 SYBR Green I Master (Roche Applied Sciences), 1 μL each primer (10 μM), 1 μl DNA template and 8 μL nuclease-free water. The 16S rRNA primers were the same as those used in the HT-qPCR. The thermal reaction conditions were as follows: 5 min pre-incubation at 95℃, then amplification of 40 cycles at 95℃ for 15 s, annealing at 60℃ for 60 s and final extension at 72℃ for 20 s. The melting curve analysis was performed by increasing the temperature stepwise from 65℃ to 97℃ using 0.30℃/5s ramp rate with continuous fluorescence recording. The results obtained by RT-qPCR exhibited a significant correlation with those obtained by HT-qPCR ($R^2 = 0.92$, $p < 0.01$). Therefore, the absolute abundance of 16S rRNA can be used to calculate the absolute abundances of ARGs and MGEs [2].

**Table S1** Descriptions of sampling sites.

| Classification | Sampling Sites | Sample abbreviation | Description | Sampling Time | Geographic Coordinates |
| --- | --- | --- | --- | --- | --- |
| Wastewater | WWTP1-S1 | AS1 | Influent | 1 Jan., 2019, July, 2019, Aug., 2019, Sep., 2020 | 23°37'39.64"N, 114°39'28.77"E |
| | WWTP1-S2 | AS2 | Anoxic tank | | |
| | WWTP1-S3 | AS3 | Aerobic tank | | |
| | WWTP1-S4 | AS4 | Effluent of biological reactor | | |
| | WWTP1-S5 | AS5 | Effluent of SSTs | | |
| | WWTP1-S6 | AS6 | Effluent of bio-contact oxidation tank | | |
| | WWTP1-S7 | AS7 | Final effluent | | |
| | WWTP2-S1 | BS1 | Influent | 10 Jan., 2019 | 23°43'4.20"N, 114°41'54.52"E |
| | WWTP2-S2 | BS2 | Anoxic tank | | |
| | WWTP2-S3 | BS3 | Aerobic tank | | |
| | WWTP2-S4 | BS4 | Effluent of biological reactor | | |
| | WWTP2-S5 | BS5 | Effluent of SSTs | | |
| | WWTP2-S6 | BS6 | Final effluent | | |
| | WWTP3-S1 | DY1 | Influent | 20 Oct., 2020 | 29°17′31.86″N, 120°18′37.15″ |
| | WWTP3-S2 | DY2 | Effluent of SSTs | | |
| | WWTP4-S1 | CX1 | Influent | 3 Nov., 2020 | 30°08′55.38″N, 121°33′38.89″E |
| | WWTP4-S2 | CX2 | Effluent of SSTs | | |
| | WWTP5-S1 | CZ1 | Influent | 25 Nov., 2020 | 31°42′58.20″N, 120°3′50.02″E |
| | WWTP5-S2 | CZ2 | Effluent of SSTs | | |
| | WWTP6-S1 | XT1 | Influent | 26 Nov., 2020 | 36°51′18.05″N, 115°30′47.38″E |
| | WWTP6-S2 | XT2 | Effluent of SSTs | | |

| | | | | | |
|---|---|---|---|---|---|
| Riverine water | Gaopu Stream-S1 | GP1 | The Gaopu Stream is located in Heyuan City, Guangdong Province, China. | 17 July, 2019 | 23°38′18.51″N, 114°38′38.54″E |
| | Gaopu Stream-S2 | GP2 | | 1 Jan., 2019 and 17 July, 2019 | 23°38′0.29″N, 114°38′55.37″E |
| | Gaopu Stream-S3 | GP3 | | | 23°37′50.17″N, 114°39′33.00″E |
| | Gaopu Stream-S4 | GP4 | | | 23°37′24.60″N, 114°40′17.50″E |
| | Yangtze River-S1 | CJ1 | Three Gorges Reservoir | 16 Sep., 2020 | 30°51′0 .72″N, 110°59′32.28″E |
| | Yangtze River-S2 | CJ2 | Huangling Temple | 16 Sep., 2020 | 30°50′22.92″N, 111°4 ′14.52″E |
| | Yangtze River-S3 | CJ3 | Xiangxi River | 17 Sep., 2020 | 30°57′26.48″N, 110°44′59.07″E |
| | Yangtze River-S4 | CJ4 | Yichang City | 18 Sep., 2020 | 30°41′19.90″N, 111°17′20.03″E |
| | Yangtze River-S5 | CJ5 | Jianli City | 18 Sep., 2020 | 29°48′28.45″N, 112°53′49.77″E |
| | Yangtze River-S6 | CJ6 | Effluent of the Dongting Lake | 19 Sep., 2020 | 29°23′14.81″N, 113°4 ′58.51″E |
| | Yangtze River-S7 | CJ7 | Anqing City | 21 Sep., 2020 | 30°30′13.52″N, 117°3 ′18.77″E |
| | Yangtze River-S8 | CJ8 | Qingyi River | 21 Sep., 2020 | 31°19′10.50″N, 118°23′38.41″E |
| | Yangtze River-S9 | CJ9 | Nanjing City | 22 Sep., 2020 | 32°5 ′44.73″N, 118°43′49.47″E |
| | Yangtze River-S10 | CJ10 | Zhenjiang City | 22 Sep., 2020 | 32°13′53.05″N, 119°21′40.11″E |
| | Yangtze River-S11 | CJ11 | Shanghai City | 23 Sep., 2020 | 31°50′22.62″N, 121°23′36.74″E |
| | Yangtze River-S12 | CJ12 | Hanjiang River | 24 Sep., 2020 | 30°35′42.39″N, 114°8 ′54.20″E |
| Stormwater runoff | SR1 | SW1 | Residential area | Stormwater event 1 was in 2 July, 2019. Stormwater event 2 was in 19 Aug., 2019. | 23°38′16.69″N, 114°38′51.44″E |
| | SR2 | SW2 | Agricultural area | | 23°38′11.52″N, 114°38′26.40″E |
| | SR3 | SW3 | Mixed flow | | 23°38′18.55″N, 114°62′62.42″E |
| | SR4 | SW4 | Mixed flow in the industrial area of Gaoxin High-tech District in Heyuan City | | 23°37′50.26″N, 114°39′32.66″E |
| | SR5 | SW5 | Mixed flow in the downstream before into the Dongjiang River | | N 23°37′24.42″N, 114°40′17.46″E |

**Table S2** Descriptions of the six full-scale WWTPs.

| Sites | Name | Scale ($m^3$/d) | Processes | Location |
|---|---|---|---|---|
| WWTP1 | Heyuan Chengnan Domestic Wastewater Treatment Plant | 25, 000 | Tridimensional eco-biological reactor | Heyuan City, Guangdong Province, China |
| WWTP2 | Heyuan Domestic Wastewater Treatment Plant | 70, 000 | AAO | Heyuan City, Guangdong Province, China |
| WWTP3 | Dongyang Domestic Wastewater Treatment Plant | 80, 000 | UBF coupled with AAO | Dongyang City, Zhejiang Province, China |
| WWTP4 | Cixi East Wastewater Treatment Plant | 100, 000 | Hydrolytic acidification coupled with modified AAO | Cixi City, Zhejiang Province, China |
| WWTP5 | Changzhou Qishuyan Wastewater Treatment Plant | 95, 000 | AAO | Changzhou City, Jiangsu Province, China |
| WWTP6 | Linxi Jieda Wastewater Treatment Plant | 20, 000 | AO | Linxi City, Hebei Province, China |

**Table S3** Primer sets of the targeted ARGs and MGEs [3, 4].

| Gene target | My Classification | Mechanisms | Forward primer | Reverse Primer |
|---|---|---|---|---|
| **16S** | | | GGGTTGCGCTCGTTGC | ATGGYTGTCGTCAGCTCGTG |
| ***aacC2*** | Aminoglycoside | deactivate | ACGGCATTCTCGATTGCTTT | CCGAGCTTCACGTAAGCATTT |
| ***aacA_aphD*** | Aminoglycoside | deactivate | AGAGCCTTGGGAAGATGAAGTTT | TTGATCCATACCATAGACTATCTCATCA |
| ***aac(6')-II*** | Aminoglycoside | deactivate | CGACCCGACTCCGAACAA | GCACGAATCCTGCCTTCTCA |
| ***acrB*** | MDR | efflux | AGTCGGTGTTCGCCGTTAAC | CAAGGAAACGAACGCAATACC |
| ***acrF*** | MDR | efflux | GCGGCCAGGCACAAAA | TACGCTCTTCCCACGGTTTC |
| ***adeA*** | MDR | efflux | CAGTTCGAGCGCCTATTTCTG | CGCCCTGACCGACCAAT |
| ***aphA3*** | Aminoglycoside | deactivate | AAAAGCCCGAAGAGGAACTTG | CATCTTTCACAAAGATGTTGCTGTCT |
| ***ermK*** | MLSB | protection | GTTTGATATTGGCATTGTCAGAGAAA | ACCATTGCCGAGTCCACTTT |
| ***multidrug resistance*** | MDR | efflux | AATTTTGCCGATTATTGCTGAAA | GATTGTCATCATTCGTTTATCACCAA |
| ***tet(36)*** | Tetracycline | protection | AGAATACTCAGCAGAGGTCAGTTCCT | TGGTAGGTCGATAACCCGAAAAT |
| ***erm(F)*** | MLSB | protection | CAGCTTTGGTTGAACATTTACGAA | AAATTCCTAAAATCACAACCGACAA |
| ***cfiA*** | Beta lactamase | deactivate | GCAGCGTTGCTGGACACA | GTTCGGGATAAACGTGGTGACT |
| ***Tp614*** | MGE | MGE | GGAAATCAACGGCATCCAGTT | CATCCATGCGCTTTTGTCTCT |
| ***IS613*** | MGE | MGE | AGGTTCGGACTCAATGCAACA | TTCAGCACATACCGCCTTGAT |
| ***blaACC-1*** | Beta lactamase | deactivate | CACACAGCTGATGGCTTATCTAAAA | AATAAACGCGATGGGTTCCA |
| ***blaMOX_blaCMY*** | Beta lactamase | deactivate | CTATGTCAATGTGCCGAAGCA | GGCTTGTCCTCTTTCGAATAGC |
| ***blaOCH*** | Beta lactamase | deactivate | GGCGACTTGCGCCGTAT | TTTTCTGCTCGGCCATGAG |
| ***blaPAO_PDC*** | Beta lactamase | deactivate | CGCCGTACAACCGGTGAT | GAAGTAATGCGGTTCTCCTTTCA |
| ***blaVEB*** | Beta lactamase | deactivate | CCCGATGCAAAGCGTTATG | GAAAGATTCCCTTTATCTATCTCAGACAA |
| ***bla1*** | Beta lactamase | deactivate | GCAAGTTGAAGCGAAAGAAAAGA | TACCAGTATCAATCGCATATACACCTAA |
| ***blaROB*** | Beta lactamase | deactivate | GCAAAGGCATGACGATTGC | CGCGCTGTTGTCGCTAAA |
| ***blaOXY-2*** | Beta lactamase | deactivate | CGTTCAGGCGGCAGGTT | GCCGCGATATAAGATTTGAGAATT |

| | | | | |
|---|---|---|---|---|
| ***blaPSE*** | Beta lactamase | deactivate | TTGTGACCTATTCCCCTGTAATAGAA | TGCGAAGCACGCATCATC |
| ***cphA*** | Beta lactamase | deactivate | GCGAGCTGCACAAGCTGAT | CGGCCCAGTCGCTCTTC |
| ***bla-L1*** | Beta lactamase | deactivate | CACCGGGTTACCAGCTGAAG | GCGAAGCTGCGCTTGTAGTC |
| ***sat4*** | Aminoglycoside | deactivate | GAATGGGCAAAGCATAAAAACTTG | CCGATTTTGAAACCACAATTATGATA |
| ***catB3*** | Amphenicol | | GCACTCGATGCCTTCCAAAA | AGAGCCGATCCAAACGTCAT |
| ***catB8*** | Amphenicol | | CACTCGACGCCTTCCAAAG | CCGAGCCTATCCAGACATCATT |
| ***ceoA*** | Amphenicol | efflux | ATCAACACGGACCAGGACAAG | GGAAAGTCCGCTCACGATGA |
| ***tet(32)*** | Tetracycline | protection | CCATTACTTCGGACAACGGTAGA | CAATCTCTGTGAGGGCATTTAACA |
| ***cmr*** | MDR | efflux | CGGCATCGTCAGTGGAATT | CGGTTCCGAAAAAGATGGAA |
| ***dfrA1*** | Other | protection | GGAATGGCCCTGATATTCCA | AGTCTTGCGTCCAACCAACAG |
| ***dfrA12*** | Other | protection | CCTCTACCGAACCGTCACACA | GCGACAGCGTTGAAACAACTAC |
| ***acrA*** | MDR | efflux | GGTCTATCACCCTACGCGCTATC | GCGCGCACGAACATACC |
| ***emrD*** | MDR | efflux | CTCAGCAGTATGGTGGTAAGCATT | ACCAGGCGCCGAAGAAC |
| ***tetU*** | Tetracycline | unknown | GTGGCAAAGCAACGGATTG | TGCGGGCTTGCAAAACTATC |
| ***vanC*** | Vancomycin | protection | CCTGCCACAATCGATCGTT | CGGCTTCATTCGGCTTGATA |
| ***lmrA*** | MLSB | efflux | TTCAGATGCAATGGCGTTTG | ATAATCGGGAACATAATGAGCATAACTAC |
| ***nisB*** | Other | deactivate | GGGAGAGTTGCCGATGTTGTA | AGCCACTCGTTAAAGGGCAAT |
| ***mdtE_yhiU*** | MDR | efflux | CGTCGGCGCACTCGTT | TCCAGACGTTGTACGGTAACCA |
| ***mexA*** | MDR | efflux | AGGACAACGCTATGCAACGAA | CCGGAAAGGGCCGAAAT |
| ***erm(36)*** | MLSB | protection | GGCGGACCGACTTGCAT | TCTGCGTTGACGACGGTTAC |
| ***aac(6')-Ib*** | Aminoglycoside | deactivate | CGTCGCCGAGCAACTTG | CGGTACCTTGCCTCTCAAACC |
| ***aadA2-1*** | Aminoglycoside | deactivate | ACGGCTCCGCAGTGGAT | GGCCACAGTAACCAACAAATCA |
| ***aadA5*** | Aminoglycoside | deactivate | ATCACGATCTTGCGATTTTGCT | CTGCGGATGGGCCTAGAAG |
| ***aadA_99*** | Aminoglycoside | deactivate | GTTGTGCACGACGACATCATT | GGCTCGAAGATACCTGCAAGAA |
| ***acrR*** | MDR | regulator | GCGCTGGAGACACGACAAC | GCCTTGCTGCGAGAACAAA |

| | | | | |
|---|---|---|---|---|
| ***aph(2')-Id*** | Aminoglycoside | deactivate | TGAGCAGTATCATAAGTTGAGTGAAAAG | GACAGAACAATCAATCTCTATGGAATG |
| ***cfxA*** | Beta lactamase | deactivate | TCATTCCTCGTTCAAGTTTTCAGA | TGCAGCACCAAGAGGAGATGT |
| ***cepA*** | Beta lactamase | deactivate | AGTTGCGCAGAACAGTCCTCTT | TCGTATCTTGCCCGTCGATAAT |
| ***blaCMY*** | Beta lactamase | deactivate | AAAGCCTCATGGGTGCATAAA | ATAGCTTTTGTTTGCCAGCATCA |
| ***ampC_blaDHA*** | Beta lactamase | deactivate | TGGCCGCAGCAGAAAGA | CCGTTTTATGCACCCAGGAA |
| ***blaGES*** | Beta lactamase | deactivate | GCAATGTGCTCAACGTTCAAG | GTGCCTGAGTCAATTCTTTCAAAG |
| ***blaSFO*** | Beta lactamase | deactivate | CCGCCGCCATCCAGTA | GGGCCGCCAAGATGCT |
| ***blaTLA*** | Beta lactamase | deactivate | ACACTTTGCCATTGCTGTTTATGT | TGCAAATTTCGGCAATAATCTTT |
| ***blaZ*** | Beta lactamase | deactivate | GGAGATAAAGTAACAAATCCAGTTAGATATGA | TGCTTAATTTTCCATTTGCGATAAG |
| ***qacF_H*** | MDR | efflux | TCGCAACATCCGCATTAAAA | ATGGATTTCAGAACCAGAGAAAGAAA |
| ***cmlA1*** | Amphenicol | | TAGGAAGCATCGGAACGTTGAT | CAGACCGAGCACGACTGTTG |
| ***cmx(A)*** | Amphenicol | efflux | GCGATCGCCATCCTCTGT | TCGACACGGAGCCTTGGT |
| ***catA1*** | Amphenicol | | GGGTGAGTTTCACCAGTTTTGATT | CACCTTGTCGCCTTGCGTATA |
| ***sul2*** | Sulfonamide | protection | TCATCTGCCAAACTCGTCGTTA | GTCAAAGAACGCCGCAATGT |
| ***ermT*** | MLSB | protection | GTTCACTAGCACTATTTTTAATGACAGAAGT | GAAGGGTGTCTTTTTAATACAATTAACGA |
| ***msr(C)*** | MLSB | protection | TCAGACCGGATCGGTTGTC | CCTATTTTTTGGAGTCTTCTCTCTAATGTT |
| ***mphB*** | MLSB | deactivate | CGCAGCGCTTGATCTTGTAG | TTACTGCATCCATACGCTGCTT |
| ***blaVIM*** | Beta lactamase | deactivate | GCACTTCTCGCGGAGATTG | CGACGGTGATGCGTACGTT |
| ***msr(A)*** | MLSB | efflux | CTGCTAACACAAGTACGATTCCAAAT | TCAAGTAAAGTTGTCTTACCTACACCATT |
| ***aadD*** | Aminoglycoside | deactivate | CCGACAACATTTCTACCATCCTT | ACCGAAGCGCTCGTCGTATA |
| ***nimE*** | MDR | deactivate | TGCGCCAAGATAGGGCATA | GTCGTGAATTCGGCAGGTTTA |
| ***Pbp5*** | Beta lactamase | protection | GGCGAACTTCTAATTAATCCTATCCA | CGCCGATGACATTCTTCTTATCTT |
| ***pbp*** | Beta lactamase | protection | CCGGTGCCATTGGTTTAGA | AAAATAGCCGCCCCAAGATT |
| ***mecA*** | Beta lactamase | protection | GGTTACGGACAAGGTGAAATACTGAT | TGTCTTTTAATAAGTGAGGTGCGTTAATA |
| ***emrB_qacA*** | MDR | efflux | CTTTTCTCTAACCGTACATTATCTACGATAAA | AGAACGTAGCGACTGATAAAATGCT |

| | | | | |
|---|---|---|---|---|
| ***blaCTX-M*** | Beta lactamase | deactivate | GCGATAACGTGGCGATGAAT | GTCGAGACGGAACGTTTCGT |
| ***aadA2-2*** | Aminoglycoside | deactivate | CAATGACATTCTTGCGGGTATC | GACCTACCAAGGCAACGCTATG |
| ***aadA9*** | Aminoglycoside | deactivate | CGCGGCAAGCCTATCTTG | CAAATCAGCGACCGCAGACT |
| ***aphA1*** | Aminoglycoside | deactivate | TGAACAAGTCTGGAAAGAAATGCA | CCTATTAATTTCCCCTCGTCAAAAA |
| ***aadE*** | Aminoglycoside | deactivate | TACCTTATTGCCCTTGGAAGAGTTA | GGAACTATGTCCCTTTTAATTCTACAATCT |
| ***str*** | Aminoglycoside | deactivate | AATGAGTTTTGGAGTGTCTCAACGTA | AATCAAAACCCCTATTAAAGCCAAT |
| ***strA*** | Aminoglycoside | deactivate | CCGGTGGCATTTGAGAAAAA | GTGGCTCAACCTGCGAAAAG |
| ***strB*** | Sulfonamide | protection | GCTCGGTCGTGAGAACAATCT | CAATTTCGGTCGCCTGGTAGT |
| ***tetA*** | Tetracycline | efflux | CTCACCAGCCTGACCTCGAT | CACGTTGTTATAGAAGCCGCATAG |
| ***tetB*** | Tetracycline | efflux | AGTGCGCTTTGGATGCTGTA | AGCCCCAGTAGCTCCTGTGA |
| ***tetK*** | Tetracycline | efflux | CAGCAGTCATTGGAAAATTATCTGATTATA | CCTTGTACTAACCTACCAAAAATCAAAATA |
| ***tetQ*** | Tetracycline | protection | CGCCTCAGAAGTAAGTTCATACACTAAG | TCGTTCATGCGGATATTATCAGAAT |
| ***tetH*** | Tetracycline | efflux | TTTGGGTCATCTTACCAGCATTAA | TTGCGCATTATCATCGACAGA |
| ***tetW*** | Tetracycline | protection | ATGAACATTCCCACCGTTATCTTT | ATATCGGCGGAGAGCTTATCC |
| ***tetO*** | Tetracycline | protection | CAACATTAACGGAAAGTTTATTGTATACCA | TTGACGCTCCAAATTCATTGTATC |
| ***tetL*** | Tetracycline | efflux | ATGGTTGTAGTTGCGCGCTATAT | ATCGCTGGACCGACTCCTT |
| ***tetX*** | Tetracycline | deactivate | AAATTTGTTACCGACACGGAAGTT | CATAGCTGAAAAAATCCAGGACAGTT |
| ***tetC*** | Tetracycline | efflux | ACTGGTAAGGTAAACGCCATTGTC | ATGCATAAACCAGCCATTGAGTAAG |
| ***tetS*** | Tetracycline | protection | TTAAGGACAAACTTTCTGACGACATC | TGTCTCCCATTGTTCTGGTTCA |
| ***tnpA-01*** | MGE | MGE | GCCGCACTGTCGATTTTTATC | GCGGGATCTGCCACTTCTT |
| ***tnpA-02*** | MGE | MGE | CCGATCACGGAAAGCTCAAG | GGCTCGCATGACTTCGAATC |
| ***tnpA-03*** | MGE | MGE | GGGCGGGTCGATTGAAA | GTGGGCGGGATCTGCTT |
| ***tnpA-04*** | MGE | MGE | CATCATCGGACGGACAGAATT | GTCGGAGATGTGGGTGTAGAAAGT |
| ***tnpA-05*** | MGE | MGE | GAAACCGATGCTACAATATCCAATTT | CAGCACCGTTTGCAGTGTAAG |
| ***tnpA-06*** | MGE | MGE | TGCAGATGGTTTAACCTTGGATATTT | TCGGTTCATCAAACTGCTTCAC |

| | | | | |
|---|---|---|---|---|
| ***tnpA-07*** | MGE | MGE | AATTGATGCGGACGGCTTAA | TCACCAAACTGTTTATGGAGTCGTT |
| ***folA*** | Sulfonamide | protection | CGAGCAGTTCCTGCCAAAG | CCCAGTCATCCGGTTCATAATC |
| ***ermX*** | MLSB | protection | GCTCAGTGGTCCCCATGGT | ATCCCCCCGTCAACGTTT |
| ***VanB*** | Vancomycin | protection | TTGTCGGCGAAGTGGATCA | AGCCTTTTTCCGGCTCGTT |
| ***vanD*** | Vancomycin | protection | CAGAGGAACATAATGTTTCGATAAAATCT | GCCGGATTTTGTGATTCCAA |
| ***vanHD*** | Vancomycin | protection | GTGGCCGATTATACCGTCATG | CGCAGGTCATTCAGGCAAT |
| ***vanHB*** | Vancomycin | protection | GAGGTTTCCGAGGCGACAA | CTCTCGGCGGCAGTCGTAT |
| ***vanRA*** | Vancomycin | protection | CCCTTACTCCCACCGAGTTTT | TTCGTCGCCCCATATCTCAT |
| ***vanSA*** | Vancomycin | protection | CGCGTCATGCTTTCAAAATTC | TCCGCAGAAAGCTCAATTTGTT |
| ***vanWB*** | Vancomycin | protection | CGGACAAAGATACCCCCTATAAAG | AAATAGTAAATTGCTCATCTGGCACAT |
| ***vanXB*** | Vancomycin | protection | AGGCACAAAATCGAAGATGCTT | GGGTATGGCTCATCAATCAACTT |
| ***vgaB*** | MLSB | efflux | TAAAAGAGAATAAGGCGCAAGGA | TGTTTAGTAGCATGTTGCATTTTCC |
| ***pica*** | MLSB | protection | GCAATCGAGGCGGTGTTC | TTGCCGCAGCCAATTCA |
| ***mtrE*** | MDR | efflux | CGATGTGTCGTTTTGGAAGGT | CCTGCACCATGATTCCTCAATA |
| ***oprD*** | MDR | efflux | ATGAAGTGGAGCGCCATTG | GGCCACGGCGAACTGA |
| ***penA*** | Beta lactamase | protection | AGACGGTAACGTATAACTTTTTGAAAGA | GCGTGTAGCCGGCAATG |
| ***pmrA*** | Other | deactivate | TTTGCAGGTTTTGTTCCTAATGC | GCAGAGCCTGATTTCTCCTTTG |
| ***ttgA*** | MDR | efflux | ACGCCAATGCCAAACGATT | GTCACGGCGCAGCTTGA |
| ***ttgB*** | MDR | efflux | TCGCCCTGGATGTACACCTT | ACCATTGCCGACATCAACAAC |
| ***mepA*** | MDR | efflux | ATCGGTCGCTCTTCGTTCAC | ATAAATAGGATCGAGCTGCTGGAT |
| ***mexE*** | MDR | efflux | GGTCAGCACCGACAAGGTCTAC | AGCTCGACGTACTTGAGGAACAC |
| ***qnrA*** | Fluoroquinolone | protection | AGGATTTCTCACGCCAGGATT | CCGCTTTCAATGAAACTGCAA |
| ***lnuA*** | MLSB | deactivate | TGACGCTCAACACACTCAAAAA | TTCATGCTTAAGTTCCATACGTGAA |
| ***mtrD*** | MDR | efflux | CGGAGTCCATCGACCATTTG | ATCGTCGGCAAGGAGAATCA |
| ***vat(E)*** | MLSB | vat(E) | GACCGTCCTACCAGGCGTAA | TTGGATTGCCACCGACAATT |

| | | | | |
|---|---|---|---|---|
| ***ermY*** | MLSB | protection | TTGTCTTTGAAAGTGAAGCAACAGT | TAACGCTAGAGAACGATTTGTATTGAG |
| ***cfr*** | MLSB | protection | GCAAAATTCAGAGCAAGTTACGAA | AAAATGACTCCCAACCTGCTTTAT |
| ***sulA_folP*** | Sulfonamide | protection | CAGGCTCGTAAATTGATAGCAGAAG | CTTTCCTTGCGAATCGCTTT |
| ***ermA_ermTR*** | MLSB | protection | ACATTTTACCAAGGAACTTGTGGAA | GTGGCATGACATAAACCTTCATCA |
| ***oleC*** | MLSB | efflux | CCCGGAGTCGATGTTCGA | GCCGAAGACGTACACGAACAG |
| ***carB*** | MLSB | efflux | GGAGTGAGGCTGACCGTAGAAG | ATCGGCGAAACGCACAAA |
| ***pikR2*** | MLSB | protection | TCGTGGGCCAGGTGAAGA | TTCCCCTTGCCGGTGAA |
| ***tetE*** | Tetracycline | efflux | TTGGCGCTGTATGCAATGAT | CGACGACCTATGCGATCTGA |
| ***tetbP*** | Tetracycline | efflux | TGGGCGACAGTAGGCTTAGAA | TGACCCTACTGAAACATTAGAAATATACCT |
| ***tetT*** | Tetracycline | protection | CCATATAGAGGTTCCACCAAATCC | TGACCCTATTGGTAGTGGTTCTATTG |
| ***tolC*** | MDR | efflux | GGCCGAGAACCTGATGCA | AGACTTACGCAATTCCGGGTTA |
| ***vanRB*** | Vancomycin | protection | GCCCTGTCGGATGACGAA | TTACATAGTCGTCTGCCTCTGCAT |
| ***vanRC*** | Vancomycin | protection | TGCGGGAAAAACTGAACGA | CCCCCCATACGGTTTTGATTA |
| ***vanRC4*** | Vancomycin | protection | AGTGCTTTGGCTTATCTCGAAAA | TCCGGCAGCATCACATCTAA |
| ***vanRD*** | Vancomycin | protection | TTATAATGGCAAGGATGCACTAAAGT | CGTCTACATCCGGAAGCATGA |
| ***vanSC*** | Vancomycin | protection | ATCAACTGCGGGAGAAAAGTCT | TCCGCTGTTCCGCTTCTT |
| ***vanTE*** | Vancomycin | protection | GTGGTGCCAAGGAAGTTGCT | CGTAGCCACCGCAAAAAAAT |
| ***vanTC*** | Vancomycin | protection | ACAGTTGCCGCTGGTGAAG | CGTGGCTGGTCGATCAAAA |
| ***vanTG*** | Vancomycin | protection | CGTGTAGCCGTTCCGTTCTT | CGGCATTACAGGTATATCTGGAAA |
| ***vanYB*** | Vancomycin | protection | GGCTAAAGCGGAAGCAGAAA | GATATCCACAGCAAGACCAAGCT |
| ***vanYD*** | Vancomycin | protection | AAGGCGATACCCTGACTGTCA | ATTGCCGGACGGAAGCA |
| ***imp-marko*** | Beta lactamase | deactivate | GGAATAGAGTGGCTTAATTC | GGTTTAACAAAACAACCACC |
| ***qnrB-bob_resign*** | Amphenicol | efflux | GCGACGTTCAGTGGTTCAGA | GCTGCTCGCCAGTCGAA |
| ***merA-marko*** | MDR | unknown | GTGCCGTCCAAGATCATG | GGTGGAAGTCCAGTAGGGTGA |
| ***int1*** | MGE | MGE | CGAAGTCGAGGCATTTCTGTC | GCCTTCCAGAAAACCGAGGA |

| | | | | |
|---|---|---|---|---|
| ***intl2*** | MGE | MGE | TGCTTTTCCCACCCTTACC | GACGGCTACCCTCTGTTATCTC |
| ***IncN_rep*** | MGE | MGE | AGTTCACCACCTACTCGCTCCG | CAAGTTCTTCTGTTGGGATTCCG |
| ***IncN_oriT*** | MGE | MGE | TTGGGCTTCATAGTACCC | GTGTGATAGCGTGATTTATGC |
| ***IncP_oriT*** | MGE | MGE | CAGCCTCGCAGAGCAGGAT | CAGCCGGGCAGGATAGGTGAAGT |
| ***IncQ_oriT*** | MGE | MGE | TTCGCGCTCGTTGTTCTTCGAGC | GCCGTTAGGCCAGTTTCTCG |
| ***IncW_trwAB*** | MGE | MGE | AGCGTATGAAGCCCGTGAAGGG | AAAGATAAGCGGCAGGACAATAACG |
| ***qacH_351*** | MDR | efflux | GTCGGTGTTGCTTATGCAGTCT | CAACCAGGCAATGGCTGTAA |
| ***marR*** | MDR | regulator | GCTGTTGATGACATTGCTCACA | CGGCGTACTGGTGAAGCTAAC |
| ***trfa*** | MGE | MGE | ACGAAGAAATGGTTGTCCTGTTC | CGTCAGCTTGCGGTACTTCTC |
| ***NDM new*** | Beta lactamase | deactivate | GGCCACACCAGTGACAATATCA | CAGGCAGCCACCAAAAGC |
| ***sul1 NEW*** | Sulfonamide | protection | GCCGATGAGATCAGACGTATTG | CGCATAGCGCTGGGTTTC |
| ***orf37-IS26*** | MGE | MGE | GCCGGGTTGTGCAAATAGAC | TGGCAATCTGTCGCTGCTG |
| ***orf39-IS26*** | MGE | MGE | GCGCGTCGAGCATCAATAG | CAGTTGTGCTGCTGGTGGTC |
| ***ISPps1-pseud*** | MGE | MGE | CACACTGCAAAAACGCATCCT | TGTCTTTGGCGTCACAGTTCTC |
| ***ISSm2-Xanthob*** | MGE | MGE | TGGATCGACCGGTTCCAT | GCTGACCGAGCTGTCCATGT |
| ***ISAba3-Acineto*** | MGE | MGE | TCAGAGGCAGCGGTATACGA | GGTTGATTCAGTTAAAGTACGTAAAACTTT |
| ***ISEfm1-Entero*** | MGE | MGE | AGGTGTCCATGACGTGAAAGTG | TCCTTTGTCCCCTAGGATATTGG |
| ***mexB*** | MDR | efflux | CTGGAGATCGACGACGAGAAG | GAAATCGTTGACGTAGCTGGAA |
| ***cmlA5*** | Amphenicol | efflux | GCGCTCTTCGAGGATTCG | CCGCCCAAGCAGAAGTAGAC |
| ***IS1111*** | MGE | MGE | GTCTTAAGGTGGGCTGCGTG | CCCCGAATCTCATTGATCAGC |
| ***PAMBL-1-F_377*** | MGE | MGE | CAGGCTCTTAATGTGATA | TTATGCTCAATACTCGTG |
| ***pAKD1-IncP-1β*** | MGE | MGE | GGTAAGATTACCGATAAACT | GTTCGTGAAGAAGATGTA |
| ***pBS228-IncP-1α*** | MGE | MGE | CAATCCATCGACAATCAC | GACAATCAGCTACTTCAC |
| ***IS1133*** | MGE | MGE | GCAGCGTCGGGTTGGA | ACGCGTTCGAACAACTGTAATG |
| ***TN5*** | MGE | MGE | CAGCATAAAAAATCCCGACAACA | CCCCGCAACAGACATACGT |

| | | | | |
|---|---|---|---|---|
| ***aac3ia*** | Aminoglycoside | | ACGTTCTGCCAAAGTTTGAG | ACTGCCGGATCGTCAC |
| ***aph4ib*** | Aminoglycoside | | GGGAACACCGTGCTCACC | GTTGGTCCCGTGCAGGTC |
| ***aph3via*** | Aminoglycoside | | TCTCATGGCGATATCACGGATAG | TTTCCTCCGATGCATCCTCTC |
| ***aph6ic*** | Aminoglycoside | | CACGACAACGTGCTCGAC | CCGTCTTCGGCGAACCA |
| ***ArmA*** | Aminoglycoside | | TCTTCGACGAATGAAAGAGTCG | GCTAATGGATTGAAGCCACAACC |
| ***spcN*** | Aminoglycoside | deactivate | GCTATGTGCTGGTGGACTGG | GGAACCACTCGACGAACTCG |
| ***spec_aph*** | Aminoglycoside | deactivate | GGTGCTGATATGAATGCCTTTGG | CATTGGGCGCATCAATAAATGG |
| ***aac(3)-ib*** | Aminoglycoside | | CAGCGAGACGTTCATCGC | CACGCTTCAGGTGGCTAATC |
| ***aac(3)-id_ie*** | Aminoglycoside | | AGATAGTTATGCCCGCAACAAG | ACGCGCTGCGCCTATA |
| ***aac(3)-iid_iii_iif_iia_iie*** | Aminoglycoside | | CGATGGTCGCGGTTGGTC | TCGGCGTAGTGCAATGCG |
| ***aac(3)-xa*** | Aminoglycoside | | GCAAGCGGTTCGTGACGTA | TCAGGTGCTCCTCGATCCAG |
| ***Aac6-Aph2*** | Aminoglycoside | | CCAAGAGCAATAAGGGCATACCAA | GCCACACTATCATAACCACTACCG |
| ***aac(6)-ig*** | Aminoglycoside | | GCGATGTTAGAAGCCTCAATTCG | CACACTTCGGCCTGTCGAA |
| ***aac(6)-iic*** | Aminoglycoside | | CAGTCTTTGGCTAATCCATCACAG | AACGAACCCGGCCTTCTC |
| ***aac(6)-ij*** | Aminoglycoside | | ATGCCTGTATCTGAATCCCTGATG | GGCAATCGCTTGTTGAGTATCTG |
| ***aac(6)-im*** | Aminoglycoside | | CGTGAGCATTATACAGAGCAATGG | CCATTTCCGTTCGTAGATATTGGC |
| ***aac(6)-ir*** | Aminoglycoside | | GCTATAACGATCAGCAGCAAGC | CGCGATGCATGGCATGAC |
| ***aac(6)-is_iu_ix*** | Aminoglycoside | | AAGCTTACTCTGGCCTGATCATG | TGCCTGAACGTCGATATTCAGG |
| ***aac(6)-iv_ih*** | Aminoglycoside | | TTGGCTTATACCGACACCCA | CCCGTTGCGATACCTGAAC |
| ***aac(6)-iw*** | Aminoglycoside | | TGCGTCAGTTACTTACACGAAC | CCTGATGCATTGCATGACTGA |
| ***aac(6)-iz*** | Aminoglycoside | | TGCGCCATGACTACGTGAAC | GACTGTCCGAAGCCAGTTCG |
| ***aacA43*** | Aminoglycoside | | CTTGGCCTACATTAGATTCAGCTC | GCTCTCAATCTTTGATAGGAGCAG |
| ***aadA6*** | Aminoglycoside | | CCATCGAGCGTCATCTGGAA | CCCGTCTGGCCGGATAAC |
| ***aadA7*** | Aminoglycoside | | CACTCCGCGCCTTGGA | TGTGGCGGGCTCGAAG |
| ***aadA10*** | Aminoglycoside | | ACAGGCACTCAACGTCATCG | CGCGGAGAACTCTGCTTTGA |

| | | | | |
|---|---|---|---|---|
| ***aadA16*** | Aminoglycoside | | ACGGTGGCCTGAAGCC | GAATTGCAGTTCCCGTCTGG |
| ***aadA17*** | Aminoglycoside | | TGTACGGCTCCGCAGTG | CACGGAATGATGTCGTCGTG |
| ***aadB*** | Aminoglycoside | | CCTGCTTGGTGGGCAGAC | CGGCACGCAAGACCTCAA |
| ***ant4-ib*** | Aminoglycoside | | GATGGCCGCTGACACATG | TCAACATTGCGCCATAGTGG |
| ***ant6-ia*** | Aminoglycoside | | TCGCCATGAGCTGCTGA | CCTATCATACTCCGGATAGGCATA |
| ***aph3-ib*** | Aminoglycoside | | AACAGGTTTGGGAGGCGATG | CGCAACAAGCCTCTCCTGAA |
| ***aph3-viia*** | Aminoglycoside | | CTCTCTCATGGAGATATGAGCGCTA | AATCCGGTTCAAGTCCCAACATG |
| ***aph4-ia*** | Aminoglycoside | | CGCTCCCGATTCCGGAA | CACAGTTTGCCAGTGATACACA |
| ***aph(3'')-ia*** | Aminoglycoside | | TAACAGCGATCGCGTATTTCG | TCCGACTCGTCCAACATCAATA |
| ***apmA*** | Aminoglycoside | | GGCGCACATGCATTCATCA | CTATACTCCAGTCCCACCATTTGA |
| ***aph_viii*** | Aminoglycoside | | TCGGTATCCCGGTTGTGAG | ACACGAGGTACGGGAATCC |
| ***acc3-iva*** | Aminoglycoside | deactivate | CCAACACGACGCTGCATC | GCTGTCGCCACAATGTCG |
| ***tet40*** | Tetracycline | | CTGTCCGTGCGCAATATATCC | GGATATATTGCGCACGGACAG |
| ***tetD*** | Tetracycline | efflux | AATTGCACTGCCTGCATTGC{EndPos:952} | GACAGATTGCCAGCAGCAGA{EndPos:1127} |
| ***tetPB*** | Tetracycline | efflux | TGGCAAGACGAGTTTGACTGA | GATCGCTCCACTTCAGCGATAA |
| ***tet39*** | Tetracycline | | TATAGCGGGTCCGGTAATAGGTG | CCATAACGATCCTGCCCATAGATAAC |
| ***tetG_F*** | Tetracycline | efflux | TCGCGTTCCTGCTTGCC | CCGCGAGCGACAAACCA |
| ***tetR*** | Tetracycline | regulator | CCGTCAATGCGCTGATGAC | GCCAATCCATCGACAATCACC |
| ***dfra14*** | Trimethoprim | | CGGATCATGTCATTGTTTCAGG | ATGTTAGAGGCGAAGTCTTGG |
| ***dfra17*** | Trimethoprim | | CGGGAACGGCCCTGATATTCC | CGTGTTGCGACCGCATACTTTC |
| ***dfra7*** | Trimethoprim | | GTAATCGGTAGTGGTCCTGA | ATCAGGACCACTACCGATTAC |
| ***dfra21*** | Trimethoprim | | TTGTTTCAACGCTGTCGCA | GGTTTCGGTTGAGACAAGCTC |
| ***dfra5*** | Trimethoprim | | CCATGGAGTGCCAAAGGTG | CACCTTTGGCACTCCATGG |
| ***dfrA8*** | Trimethoprim | | GGTCGCACCTGCATCGTTA | AGCGCCACCAATGACGTAG |
| ***dfrA10*** | Trimethoprim | | CTTCAACTATCACAGAGCACGAAG | TCTACCGGTACATACACATCAGC |

| | | | | |
|---|---|---|---|---|
| ***dfrA15*** | Trimethoprim | | AGGCCGAAAGACTTTCGAGTC | TCACCTTCTGGCTCAATGTCG |
| ***dfrA18*** | Trimethoprim | | GGAGCGAATCAAGGAGAAAGGAA | GCAATGCGTTGATCGGTATTCTC |
| ***dfrA22*** | Trimethoprim | | CAGCCGAACACGGCAAAG | CGGAGTGCGTGTACGTGA |
| ***dfrA25*** | Trimethoprim | | TCAAACTGGACAGCGGCTA | GTCGATTGTCGACACATGCA |
| ***dfrA27*** | Trimethoprim | | GCCGCTCAGGATCGGTA | GTCGAGATATGTAGCGTGTCG |
| ***dfrAB4*** | Trimethoprim | | CGGTTCGCATTCCCATCAAA | CGCAGTCATGGGATAAATCTGG |
| ***dfrC*** | Trimethoprim | | GTCGCTCACGATAAACAAAGAGTC | CCCTTCATGGTGAAATGAAGCTTG |
| ***dfrG*** | Trimethoprim | | TCAATCGGAAGAGCCTTACCTGA | TGGGCAAATACCTCATTCCATTCC |
| ***dfrK*** | Trimethoprim | | TGCTGCGATGGATAAGAACAG | CTTCCAGGTAATGCTCTTCCG |
| ***dfrBmulti*** | Trimethoprim | | ACCAAGGCAGAAGTGAAGTCA | GGTGAGCCTCAGACTCGAC |
| ***fosb*** | Other | | CTTGCAGGCCTATGGATTGC | TCTGTTCTCAAGTGTGCCAGTA |
| ***fosX*** | Other | deactivate | AGCTGGTTTGTGGATTTGCA | CCACACCGAGAGCTTTAATCCG |
| ***Arr2*** | Other | | TTGGCGATTGGTGACTTGCTAA | ATCGTCTTCGAACGGTCCTG |
| ***mcr-1*** | Other | | CACATCGACGGCGTATTCTG | CAACGAGCATACCGACATCG |
| ***sulIII*** | Sulfonamide | protection | CGCGCTCAAGGCAGATG | GGGAATGCCATCTGCCTTG |
| ***ere(A)*** | MLSB | deactivate | GATAATTCTGCTGGCGCACA | GCAGGCGTGGTCACAAC |
| ***ere(B)*** | MLSB | deactivate | TCGTATATGGCGGGCGTAGTA | GGTCCAAGATGGGTGAATGCA |
| ***erm(A)*** | MLSB | protection | TCGTTGAGAAGGGATTTGCGA | TTGCATGCTTCAAAGCCTGTC |
| ***erm(B)*** | MLSB | protection | GAACACTAGGGTTGTTCTTGCA | CTGGAACATCTGTGGTATGGC |
| ***erm(D)*** | MLSB | protection | TTTCCGGACAGCATTTGATGC | TCCACTGCCAATACCTTACCG |
| ***erm(E)*** | MLSB | | GTCACGCAGCTGGAGTTCG | CGGTGAAGCACAGCTCGAC |
| ***erm(G)*** | MLSB | | CCCTTGAATTAGTACAGAGGTG | GCAAACTCGTATTCCACGA |
| ***erm(O)*** | MLSB | | TGATGACGGCTCAGTGG | GTGCACCAGCGCCTGA |
| ***erm(Q)*** | MLSB | | TGAAAGCCATGCGTCTGAC | TTCAGCTGGCAGCTTAAGC |
| ***erm(S)*** | MLSB | | GAGTACGCCCGCAAACG | GCGTTCGATCCGGAGGA |

| | | | | |
|---|---|---|---|---|
| ***lnuB*** | MLSB | deactivate | GGATCGTTTACCAAAGGAGAAGG | AGCATAGCCTTCGTATCAGGAA |
| ***mphA*** | MLSB | deactivate | TCAGCGGGATGATCGACTG | GAGGGCGTAGAGGGCGTA |
| ***vat(A)*** | MLSB | | ATGAACGGAGCGAATCATCGG | CCATACCGATCCAAACGTCATTTC |
| ***erm(34)*** | MLSB | protection | AAAGCGGTTTACAAGCGTTTCG | GGGTGCTCTAGGGTTGTTTAGTG |
| ***erm(35)*** | MLSB | protection | CCTTCAGTCAGAACCGGCAA | GCTGATTTGACAGTTGGTGGTG |
| ***erm(42)*** | MLSB | | TGTTGAGATTGGGCCTGGA | CTAAGGGTGGGTTCTCACTATCTA |
| ***erm(F)*** | MLSB | | TCTGATGCCCGAAATGTTCAAG | TGAAGGACAATTGAACCTCCCA |
| ***lnu(F)*** | MLSB | | ATACCGGTCATTTCCACTTGGC | GCATCAGGCTGATGAGGTTCAA |
| ***lsa(C)*** | MLSB | | AAACGGCGTGAAAGTATCAGG | TTGTGGTGATGTAACGGATGC |
| ***mef(B)*** | MLSB | | CCGATAGGCTTACTTGTTGCAG | AGTCCACTTGCGGTTTCATTG |
| ***msr(D)*** | MLSB | | GGCAAGCTAGGTGTTGAGC | ATTGCTCAACACCTAGCTTGC |
| ***msr(E)*** | MLSB | | CGGCAGATGGTCTGAGCTTAAA | CGCACTCTTCCTGCATAAAGGA |
| ***vgaA*** | MLSB | efflux | GGAAGCTATAGAGGCGTTTGAATC | CCGAAGGTTCAATACTCAATCGAC |
| ***vga(A)LC*** | MLSB | | GTGAAGATGTCTCGGGTACAATTG | GAAATACCAGGATTCCCATGCAC |
| ***vatB*** | MLSB | deactivate | GCAATTGTTGCTGCGAATTCAG | GTGCTGACCAATCCCACCA |
| ***cat*** | Phenicol | | ATCGGCCAGACTGGATATCGA | CACAGCTCCAGTTGCAACAAC |
| ***catA2*** | Phenicol | | CCTGGAACCGCAGAGAACA | CGGAACTCCGGAAACTGATTAAC |
| ***catA3*** | Phenicol | | CTGATTGCTCAGGCCGTGAA | ATGAGTATGGGCAACTCAGTGC |
| ***catB2*** | Phenicol | | GCTACTATTCCGGCTATTACCATG | GGGCTCCTCGTTCATGTAGA |
| ***catB9*** | Phenicol | | CACCTTATGAAGTGGTCGGTTCA | GTCTGATGAACACAGAGACTGCA |
| ***cat(pC221)*** | Phenicol | | AATGACCGTATGCTGCAAGAAG | TTTGCCTGCTATGGCATTCTG |
| ***catP*** | Phenicol | | CCTTTGGACTGAGTGTAAGTCTGA | TAAAGCCATCGAAGGTTGACCA |
| ***catQ*** | Phenicol | | AGGTGCACTTACAGTATGACTGC | AACGTGGGAAGTTCTCGTCATAC |
| ***cmlV*** | Phenicol | | GCCCTCATCACCGTCTTCG | GGACGTTGGCGATGGAGAG |
| ***fexA*** | Phenicol | | TGGTGTGGCTGTTGCAATCTTA | CCAAGGTACAAAGCACCTTGGA |

| | | | | |
|---|---|---|---|---|
| ***floR*** | Amphenicol | | AACCCGCCCTCTGGATCA | GCCGTCGAGAAGAAGACGAA |
| ***vanG*** | Vancomycin | protection | TGTTTCGCAGAACCGTGTCAA | CCCTGCACTGTTCCATCTTCTC |
| ***vanC2_vanC3*** | Vancomycin | protection | TGACTGTCGGTGCTTGTGA | GATAGAGCAGCTGAGCTTGTTC |
| ***beta_ccra*** | Beta lactamase | | CACTGGCACGGCGATTGTA | CGGCAGCCAAACCACGATA |
| ***cefa_ampc*** | Beta lactamase | | CAGGATCTGATGTGGGAGAACTA | TCGGGAACCATTTGTTGGC |
| ***bl1acc*** | Beta lactamase | | TGTTATCCGTGATTACCTGTCTGG | CTCAGCGAGCCAACTTCAAATA |
| ***blaCTX-M-1_3_15*** | Beta lactamase | deactivate | CGTACCGAGCCGACGTTAA | CAACCCAGGAAGCAGGCA |
| ***blaSHV-11*** | Beta lactamase | deactivate | TTGACCGCTGGGAAACGG | TCCGGTCTTATCGGCGATAAAC |
| ***bl3_cpha*** | Beta lactamase | | GTAACGCCTACTGGAAGTCCA | CAGCTTCTCCTTGAGAATGCAG |
| ***blaB-11_13_14*** | Beta lactamase | | CGTGCCGGAGGTCTTGAATA | GGGATAGTAAACCTGAAACTCGGA |
| ***blaIND*** | Beta lactamase | | CGCCTGTTAAACCCAACCTGTA | CGCTCTGTCATCATGAGAGTGG |
| ***blaLEN*** | Beta lactamase | | TGTTCGCCTGTGTGTTATCTCC | GCAGCACTTTAAAGGTGCTCAC |
| ***blaOXY-1*** | Beta lactamase | deactivate | AAAGGTGACCGCATTCGC | CCAGCGTCAGCTTGCG |
| ***bla-SME*** | Beta lactamase | deactivate | GAGGAAGACTTTGATGGGAGGATTG | CGCTATATTGCAATGCAGCAGAAG |
| ***blaCARB*** | Beta lactamase | | TGATTTGAGGGATACGACAACTCC | CTGTAATACTCCGAGCACCAA |
| ***blaGOB*** | Beta lactamase | | CTTGGGCTTGAATGCTCAGGTA | TGTATGGTCGTAGTGAGCCTGA |
| ***blaHERA*** | Beta lactamase | | GGGCAACCGCATTCTGAC | GCATCTCCCACTTTATCGTCAC |
| ***blaMIR*** | Beta lactamase | | CGGTCTGCCGTTACAGGTG | AAAGACCCGCGTCGTCATG |
| ***blaFOXnew*** | Beta lactamase | deactivate | CCTACGGCTATTCGAAGGAAGATAAG | CCGGATTGGCCTGGAAGC |
| ***nonmobile_blaADC*** | Beta lactamase | | GGTATGGCTGTGGGTGTTATTCA | AGGCAAGGTTACCACTTGTATACG |
| ***nonmobile blaBEL*** | Beta lactamase | | ATGTCCATGGCACAGACTGTG | CCTGTCTTGTCACCCGTTACC |
| ***blaIMI*** | Beta lactamase | | ACATCTACACCTGCAGCAGTAG | AATCGCTTGGTACGCTAGCA |
| ***norA*** | Fluoroquinolone | | ATCGCCGTTTGGTGGTACG | TCCACCAATCCCTGGTCCTAAA |
| ***qepA_1_2*** | Fluoroquinolone | | GGGCATCGCGCTGTTC | GCGCATCGGTGAAGCC |
| ***QnrB4*** | Fluoroquinolone | | TCACCACCCGCACCTG | GGATATCTAAATCGCCCAGTTCC |

| | | | | |
|---|---|---|---|---|
| ***QnrS1_S3_S5*** | Fluoroquinolone | | CCACTTTGATGTCGCAGATCTTC | CCCTCTCCATATTGGCATAGGAAA |
| ***QnrVC1_VC3_VC6*** | Fluoroquinolone | | CTCACATCAGGACTTGCAAGAA | ATGAAGCATCTCGAAGATCAGC |
| ***QnrVC4_VC5_VC7*** | Fluoroquinolone | | TTCCTTTAAACGGGCAAACCTC | CGATACCTGATTCATGAAGCTAGC |
| ***mdth*** | MDR | | ATGCTGGCTGTACAAGTGATG | CACTCCAGCGGGCGATA |
| ***cefa_qacelta*** | MDR | | TAGTTGGCGAAGTAATCGCAAC | TGCGATGCCATAACCGATTATG |
| ***mdtg*** | MDR | efflux | TTCCAGCCGGTCAGCAA | GACATCTCCCGCGAGTTCG |
| ***pcoA*** | MDR | | TGGCGTATGGAGTTTCAATGC | GAATAATGCCGTGCCAGTGAA |
| ***silE*** | MDR | | GGTGGAAAGTCATCAGAGGATGA | CAAAGCCCAGCAAGGATGC |
| ***arsA*** | MDR | | CAGGTCAGCCGCATCAACC | GCCTGAAACACGGCAATTTCTTC |
| ***qacA_B*** | MDR | efflux | AAGGGCCACTGCATTAGCTG | CCAGTCCAATCATGCCTGCA |
| ***bacA*** | Other | deactivate | ATCCGCGGCACCCTGA | CCTGCTTGATGGACTTGATGAAGA |
| ***aac(6')I1*** | Aminoglycoside | deactivate | GGGAATTATCGGAATAGCTCTTGG | TTGGGCTGTTCTTCCTAGCTAA |
| ***aac(6')-Iy*** | Aminoglycoside | deactivate | GCCTCAATCCGCCACGATTA | ACGCGCTCTGTTTCCTCAAA |
| ***aph6ia*** | Aminoglycoside | deactivate | CGCTGGGAGCTGAAGAGG | AGCATCGTGCTGCTCTCC |
| ***bexA_norM*** | MDR | efflux | TCGGGCATCCCGTTTATGATC | GTAGGCTGCGCATAATACCCA |
| ***ampC*** | Beta lactamase | deactivate | CTGGCGCATACCTGGATTAC | GCCAGTTCAGCATCTCCCA |
| ***blaOXA10*** | Beta lactamase | deactivate | CGACCGAGTATGTACCTGCTTC | TCAAGTCCAATACGACGAGCTA |
| ***tetPA*** | Tetracycline | efflux | GGAAACCTTAGTTCAGTGACTTGG | CCCATTTAACCACGCACTGAA |
| ***blaIMIR*** | Beta lactamase | regulator | AGCCGGACTAGAGCTTCATG | GGCAGAACTCATCATCTGCAAA |
| ***mdtA*** | MDR | efflux | ACAAGCCCAGGGCCAAC | CCTTAATGGTGCCTTCGGTTTC |
| ***aac3-Via*** | Aminoglycoside | deactivate | GTGTCCGTCGCCAAGGA | GGTGACGGCCTTGTCGA |
| ***mefA*** | MLSB | efflux | TAATTATCGCAGCAGCTGGTTC | GTTCCCAAACGGAGTATAAGAGTG |
| ***blaTEM*** | Beta lactamase | deactivate | CGCCGCATACACTATTCTCAG | GCTTCATTCAGCTCCGGTTC |
| ***tetM*** | Tetracycline | protection | GGAGCGATTACAGAATTAGGAAGC | TCCATATGTCCTGGCGTGTC |
| ***vanA*** | Vancomycin | protection | GGGCTGTGAGGTCGGTTG | TTCAGTACAATGCGGCCGTTA |

| | | | | |
|---|---|---|---|---|
| ***vanXA*** | Vancomycin | protection | TCGTTGGGACGCTAAATATGC | GGACGGTAACCGTCCCATA |
| ***tetJ*** | Tetracycline | efflux | CAGCGCCCATACGCCATTTA | CCTACTTCAGTAGTGTGCCAAGC |
| ***blaPER*** | Beta lactamase | deactivate | GCAAATGAAGCGCAGATGC | GACCACAGTACCAGCTGGTA |
| ***tet(38)*** | Tetracycline | efflux | AAGCGACATTAGCCGGTTTAG | CTGCTCGTACTTAAGCCAAGG |
| ***lnuC*** | MLSB | deactivate | GGGTGTAGATGCTCTTCTTGGA | CTTTACCCGAAAGAGTTTCTACCG |
| ***fabK*** | Other | protection | CAGGAGCAGGAAATCCAAGC | CCAGCTTCCATTCCTTCTGC |
| ***vanSB*** | Vancomycin | protection | GAAGATAAAGAGGGAAGCGTACTC | CCGAATTGTCAGCCCTTGATAA |
| ***intl3*** | MGE | MGE | CAGGTGCTGGGCATGGA | CCTGGGCAGCATCACCA |
| ***KPC*** | Beta lactamase | deactivate | GCCGCCAATTTGTTGCTGAA | GCCGGTCGTGTTTCCCTTT |
| ***mobA*** | MGE | | GCTTCCCGTAACGAGGTAGT | CCTTGAACGGTATCAGCACG |
| ***traN*** | MGE | | GCTTGGCGGTCAGCAATT | TTAGGAATAACAATCGCTACACCTTTA |
| ***tra-A*** | MGE | | AAGTGTTCAGGGTGCTTCTGCGC | GTCATGTACATGATGACCAAAA |
| ***trb-C*** | MGE | | CGGYATWCCGSCSACRCTGCG | GCCACCTGYSBGCAGTCMCC |
| ***ISCR1*** | MGE | | ATGGTTTCATGCGGGTT | CTGAGGGTGTGAGCGAG |
| ***copA*** | MDR | | TGCACCTGACVGGSCAYAT | GVACTTCRCGGAACATRCC |
| ***23s -Enterococci*** | taxanomic | 23S rDNA | AGAAATTCCAAACGAACTTG | CAGTGCTCTACCTCCATCATT |
| ***ecfX-P. aeruginosa*** | taxanomic | ecfX | AGCGTTCGTCCTGCACAAGT | TCCACCATGCTCAGGGAGAT |
| ***mecA-Staphylococci*** | taxanomic | mecA | CGCAACGTTCAATTTAATTTTGTTAA | TGGTCTTTCTGCATTCCTGGA |
| ***gltA-K. pneumoniae*** | taxanomic | gltA | ACGGCCGAATATGACGAATTC | AGAGTGATCTGCTCATGAA |
| ***ompA-A. baumannii?*** | taxanomic | ompA | TCTTGGTGGTCACTTGAAGC | ACTCTTGTGGTTGTGGAGCA |
| ***czcA*** | MDR | | GCCTTGTTCATCGGCGAAC | GGCAATGTCGCCTTCGTTC |
| ***optrA*** | Phenicol | | GGTGGATGAAGTCCGTACGG | AGGTTAGACCTCCAAGAGCCA |
| ***tet44*** | Tetracycline | | CTCATGTAGATGCAGGAAAGACG | GTAACTGCTGCCTGAATTGTGA |
| ***aph3-III*** | Aminoglycoside | | CAGAAGGCAATGTCATACCACTTG | GACAGCCGCTTAGCCGAA |
| ***ant6-ib*** | Aminoglycoside | | AGAACATCCGACAGCACGTTC | CCAACCTTCCATGAAATCATTCGC |

| | | | | |
|---|---|---|---|---|
| ***ARR-3*** | Other | | GATCGTCTTCGAACGGTCCTG | TTTGGCGATTGGTGACTTGCT |
| ***mcr-2*** | Other | | CGGCGTACTTTAAGCGTTATGATG | GCATTTGGCATACCATGCAGATAG |
| ***bla-ACT*** | Beta lactamase | | AAGCCGCTCAAGCTGGA | GCCATATCCTGCACGTTGG |
| ***aac(3)-Xa*** | Aminoglycoside | | TGTACGGCTCCGCAGTG | CACGGAATGATGTCGTCGTG |
| ***IS26*** | MGE | | ATGGATGAAACCTACGTGAAGGTC | CGGTACTTAATCTGTCGGTGTTCA |
| ***IS3*** | MGE | | CGGTCTGAGCTTCGGGAA | AGAACTGTCACTCCGGTCTG |
| ***IS256*** | MGE | | CTTGCGCATCATTGGATGATGG | AAGAACGGCTCCAATTAAGCGA |
| ***sugE*** | MDR | | CTTAGTTATTGCTGGTCTGCTGGA | GCATCGGGTTAGCGGACTC |
| ***ISEcp1*** | MGE | | CATGCTCTGCGGTCACTTC | GACGCACCTTCTTGATGACC |
| ***IS200-1*** | MGE | | CCAAATACCGAAGACAAGCGTTC | CCAAACTGCTCGTAAAGCATCAG |
| ***IS1247*** | MGE | | CGGCCGTCACTGACCAA | TCGGCAGGTTGGTGACG |
| ***IS630*** | MGE | | CCGCCACCAGTGTGATGG | TTGGCGCTGACTGGATGC |
| ***Bacteroidetes*** | taxanomic | | GGARCATGTGGTTTAATTCGATGAT | AGCTGACGACAACCATGCAG |
| ***Firmicutes*** | taxanomic | | GGAGYATGTGGTTTAATTCGAAGCA | AGCTGACGACAACCATGCAC |
| ***TN5403*** | MGE | | AAGCGAATGGCGCGAAC | CGCGCAGGGTAAACTGC |
| ***IS200-2*** | MGE | | GCACACCCGATGGAACTGTAAA | TCGGCGGGATCTCCAGAAG |
| ***IS21-ISAs29*** | MGE | | GGTCCGTCAGGCACAAGTC | GGGATCGTATCGGCAAGCC |
| ***Tn3*** | MGE | | GCTGAGGTGTTCAGCTACATCC | GCTGAGGTAGTCACAGGCATTC |
| ***IS6_257*** | MGE | | ATATCGTGCCATTGATGCAGAG | ACCATTGCTACCTTCGTTGAAG |
| ***IS6100*** | MGE | | CGCACCGGCTTGATCAGTA | CTGCCACGCTCAATACCGA |
| ***IS15DI*** | MGE | | CAATACCTTTGATGGTGGCGTAAG | CTTACGCCACCATCAAAGGTATTG |
| ***IncN_korA*** | MGE | | GGAACGTTTGTAYCTTGTATTG | ACTCACTATCTTCTGTTGATTG |
| ***lncF_FIC*** | MGE | | GTGAACTGGCAGATGAGGAAGG | TTCTCCTCGTCGCCAAACTAGAT |
| ***IncI1_repI1*** | MGE | | CGAAAGCCGGACGGCAGAA | TCGTCGTTCCGCCAAGTTCGT |
| ***IncHI2-smr0018*** | MGE | | ATAATGATTCACCGGGGTAG | CTTCAGGCTATCGTTTCG |

| | | | | |
|---|---|---|---|---|
| ***IS91*** | MGE | | GGATGCCACTGCTGGTCA | ACAGTGGATACAGTATCTGCTGAG |
| ***IS5_IS1182*** | MGE | | TTCTCGAAGAATCGCCATGGC | GCTTTGGATCGCTCCAATCGA |
| ***cro*** | MGE | | AGATGTTATCGACCACTTCGGA | CCGCTTGGCGATAAGCG |
| ***EAE_05855*** | MGE | | CCCATCACCGCTGAACTGG | TGGGCGCTGCCATCTAAAC |
| ***tcrB*** | MDR | | GTGCCGGAACTCAAGTAGCA | GCACCGACTGCTGGACTTAA |
| ***terW*** | MDR | | TCAAAGAGCTACGCGAGTCATA | CCTTCCCTGTGGACTCACC |
| ***pbrT*** | MDR | | GATGCGCACTGGGCTTG | TCGGAATATGCGGAAATGCG |
| ***qnrD*** | Fluoroquinolone | | CGCTGGAATGGCACTGTGA | GCTCTCCATCCAACTTCACTCC |
| ***cadC*** | MDR | | CGCTCTGTGTCAGGATGAAGAG | CTTTCTTATGTGCTAGGGCGATCA |
| ***qnrS2*** | Fluoroquinolone | | TCCCGAGCAAACTTTGCCAA | GGTGAGTCCCTATCCAGCGA |
| ***oqxA*** | Fluoroquinolone | | GAGTCAACCTACCTCCACTATCA | GCTGCGAGTTATCCAGCAG |
| ***adeI*** | MDR | | CAGTCTGGTTTGCAGTAACCA | CACTCCTACAACAACAGGCAA |
| ***qnrB46_47_48*** | Fluoroquinolone | | CGACGTTCAGTGGTTCAGATCTC | GCCAAGCCGCTCCATGAG |

**Table S4** Sequence number and diversity indexes of the eDNA samples.

| Sample | Seq_num | Base_num | Mean_length | Min_length | Max_length | Shannon | Chao | Coverage |
|---|---|---|---|---|---|---|---|---|
| 1AS1 | 36933 | 15591785 | 422.16 | 262 | 515 | 4.25 | 791.00 | 0.994687 |
| 1AS2 | 31380 | 13361898 | 425.81 | 375 | 488 | 3.65 | 518.68 | 0.996832 |
| 1AS3 | 51858 | 22042551 | 425.06 | 375 | 467 | 3.20 | 619.51 | 0.997571 |
| 1AS4 | 35243 | 14970835 | 424.79 | 262 | 466 | 3.83 | 675.11 | 0.995852 |
| 1AS5 | 35782 | 14950939 | 417.83 | 333 | 431 | 4.08 | 919.63 | 0.99321 |
| 1AS6 | 39231 | 16720527 | 426.21 | 332 | 500 | 3.43 | 433.32 | 0.998156 |
| 7AS1 | 36383 | 15391734 | 423.05 | 278 | 432 | 3.83 | 467.00 | 0.996877 |
| 7AS2 | 34250 | 14498452 | 423.31 | 382 | 467 | 3.51 | 433.50 | 0.997104 |
| 7AS3 | 40326 | 17075628 | 423.44 | 375 | 431 | 3.73 | 623.72 | 0.995702 |
| 7AS4 | 39071 | 16679638 | 426.91 | 213 | 434 | 2.22 | 491.04 | 0.996911 |
| 7AS5 | 41414 | 17323468 | 418.30 | 397 | 431 | 2.89 | 363.50 | 0.99803 |
| 7AS6 | 34923 | 14953130 | 428.17 | 377 | 483 | 1.78 | 226.00 | 0.998238 |
| 7AS7 | 43351 | 18527169 | 427.38 | 256 | 432 | 2.13 | 262.23 | 0.99863 |
| 8AS1 | 39273 | 16544370 | 421.27 | 317 | 431 | 2.91 | 394.29 | 0.997371 |
| 8AS2 | 48524 | 20376476 | 419.93 | 265 | 469 | 3.40 | 569.09 | 0.997153 |
| 8AS3 | 44343 | 19012198 | 428.75 | 341 | 442 | 1.89 | 275.63 | 0.99832 |
| 8AS4 | 43272 | 18513600 | 427.84 | 317 | 447 | 2.79 | 401.17 | 0.997713 |
| 8AS5 | 36839 | 15668910 | 425.33 | 252 | 431 | 3.12 | 285.50 | 0.998821 |
| 8AS6 | 37061 | 15839508 | 427.39 | 362 | 431 | 2.69 | 334.53 | 0.997536 |
| 8AS7 | 35865 | 15290411 | 426.33 | 398 | 431 | 2.78 | 316.28 | 0.997747 |
| 9AS1 | 32852 | 14088389 | 428.84 | 235 | 467 | 1.67 | 307.57 | 0.997014 |
| 9AS2 | 31927 | 13691547 | 428.84 | 251 | 467 | 0.53 | 321.67 | 0.996667 |

| | | | | | | | | |
|---|---|---|---|---|---|---|---|---|
| 9AS3 | 33397 | 14327155 | 429.00 | 240 | 450 | 2.23 | 356.75 | 0.997221 |
| 9AS4 | 41196 | 17547320 | 425.95 | 220 | 516 | 2.06 | 1117.66 | 0.990929 |
| 1BS1 | 43434 | 18467031 | 425.17 | 361 | 452 | 3.61 | 950.24 | 0.994298 |
| 1BS2 | 37499 | 15339859 | 409.07 | 374 | 507 | 4.31 | 917.58 | 0.994651 |
| 1BS3 | 30566 | 12787631 | 418.36 | 262 | 467 | 4.46 | 934.28 | 0.993554 |
| 1BS4 | 31659 | 13187467 | 416.55 | 216 | 479 | 4.01 | 523.62 | 0.997345 |
| 1BS5 | 37612 | 15398024 | 409.39 | 321 | 452 | 4.15 | 604.88 | 0.997613 |
| 1BS6 | 30713 | 12967119 | 422.20 | 371 | 452 | 3.99 | 626.01 | 0.995337 |
| CX1 | 38616 | 16563201 | 428.92 | 375 | 431 | 0.63 | 215.27 | 0.998542 |
| CX2 | 31226 | 13296198 | 425.81 | 364 | 479 | 2.55 | 418.75 | 0.996766 |
| CZ1 | 40600 | 16845821 | 414.92 | 374 | 470 | 4.94 | 1676.58 | 0.9931 |
| CZ2 | 45534 | 18571000 | 407.85 | 289 | 479 | 4.06 | 1032.27 | 0.994657 |
| DY1 | 40232 | 16993547 | 422.39 | 265 | 467 | 2.65 | 712.14 | 0.993522 |
| DY2 | 42210 | 17307776 | 410.04 | 373 | 512 | 4.84 | 1325.68 | 0.993473 |
| XT1 | 43474 | 18003484 | 414.12 | 374 | 467 | 5.46 | 1585.27 | 0.995357 |
| XT2 | 43111 | 17588360 | 407.98 | 373 | 526 | 3.01 | 811.13 | 0.995505 |
| 1GP2 | 54080 | 22981484 | 424.95 | 248 | 441 | 4.13 | 542.00 | 0.997923 |
| 1GP3 | 48253 | 20557939 | 426.04 | 262 | 431 | 3.41 | 543.15 | 0.997207 |
| 7GP1 | 49747 | 21248031 | 427.12 | 387 | 431 | 2.93 | 300.03 | 0.998705 |
| 7GP2 | 46852 | 19819359 | 423.02 | 282 | 519 | 3.75 | 489.00 | 0.997956 |
| 7GP3 | 47968 | 20279411 | 422.77 | 400 | 503 | 4.25 | 532.42 | 0.997687 |
| 7GP4 | 40732 | 17360721 | 426.22 | 306 | 504 | 3.77 | 339.36 | 0.998335 |
| CJ1 | 49474 | 21112082 | 426.73 | 392 | 455 | 2.21 | 178.75 | 0.999125 |
| CJ2 | 37316 | 15967893 | 427.91 | 329 | 431 | 2.28 | 179.40 | 0.99884 |
| CJ3 | 38842 | 16651037 | 428.69 | 334 | 431 | 2.12 | 166.91 | 0.998932 |

| | | | | | | | | |
|---|---|---|---|---|---|---|---|---|
| CJ4 | 36573 | 15670533 | 428.47 | 342 | 439 | 2.29 | 260.33 | 0.998461 |
| CJ5 | 45247 | 19316144 | 426.90 | 313 | 463 | 2.49 | 237.00 | 0.998651 |
| CJ6 | 42088 | 17921310 | 425.81 | 402 | 433 | 3.01 | 417.44 | 0.998021 |
| CJ7 | 44452 | 19052147 | 428.60 | 322 | 444 | 1.95 | 409.25 | 0.997119 |
| CJ8 | 44392 | 19034414 | 428.78 | 235 | 432 | 1.91 | 224.23 | 0.998715 |
| CJ9 | 32175 | 13741304 | 427.08 | 246 | 449 | 1.38 | 193.27 | 0.99829 |
| CJ10 | 38498 | 16300664 | 423.42 | 373 | 432 | 2.44 | 298.88 | 0.998131 |
| CJ11 | 42544 | 18180885 | 427.34 | 374 | 432 | 1.44 | 275.64 | 0.998447 |
| CJ12 | 34916 | 14926421 | 427.50 | 253 | 432 | 0.85 | 248.33 | 0.998194 |
| 7SW1 | 53775 | 22543634 | 419.22 | 332 | 431 | 3.96 | 440.50 | 0.998654 |
| 7SW2 | 53680 | 22931499 | 427.19 | 316 | 452 | 2.46 | 250.00 | 0.998774 |
| 7SW3 | 57409 | 24203675 | 421.60 | 249 | 431 | 2.84 | 279.90 | 0.99901 |
| 7SW4 | 49039 | 20644448 | 420.98 | 239 | 431 | 4.02 | 410.62 | 0.99803 |
| 7SW5 | 34433 | 14661040 | 425.78 | 314 | 447 | 3.57 | 336.59 | 0.998098 |
| 8SW1 | 46248 | 19356761 | 418.54 | 378 | 526 | 4.53 | 562.33 | 0.997933 |
| 8SW2 | 38839 | 16008615 | 412.18 | 255 | 456 | 4.27 | 538.97 | 0.99754 |
| 8SW3 | 45320 | 19236094 | 424.45 | 399 | 440 | 3.42 | 376.12 | 0.998337 |
| 8SW4 | 47346 | 19887932 | 420.06 | 392 | 519 | 4.35 | 519.69 | 0.997902 |
| 8SW5 | 46873 | 19692776 | 420.13 | 350 | 431 | 4.26 | 597.00 | 0.997287 |

**Table S5** Sequence number and diversity indexes of the iDNA samples.

| Sample | Seq_num | Base_num | Mean_length | Min_length | Max_length | Shannon | Chao | Coverage |
|---|---|---|---|---|---|---|---|---|
| 1AS1 | 45509 | 19367016 | 425.5645 | 277 | 466 | 4.584621 | 1376.031 | 0.972291 |
| 1AS2 | 54881 | 22759839 | 414.7125 | 245 | 445 | 4.462349 | 1030.561 | 0.980349 |
| 1AS3 | 41994 | 17416485 | 414.7375 | 333 | 432 | 4.768387 | 930.5938 | 0.985156 |
| 1AS4 | 46736 | 19614518 | 419.6876 | 239 | 432 | 4.806362 | 1392.636 | 0.973351 |
| 1AS5 | 53709 | 22491099 | 418.7585 | 252 | 499 | 6.094234 | 2787.381 | 0.945571 |
| 1AS6 | 45939 | 19230939 | 418.619 | 277 | 499 | 5.729137 | 1664.768 | 0.972786 |
| 7AS1 | 37572 | 15958034 | 424.7321 | 337 | 466 | 4.074409 | 1485.654 | 0.971584 |
| 7AS2 | 38168 | 15902125 | 416.635 | 226 | 432 | 5.192302 | 1446.622 | 0.972079 |
| 7AS3 | 40778 | 17008830 | 417.108 | 258 | 479 | 5.198657 | 1416.944 | 0.973987 |
| 7AS4 | 46321 | 19197463 | 414.4441 | 283 | 512 | 5.106549 | 1684.084 | 0.969605 |
| 7AS5 | 52053 | 21556437 | 414.1248 | 285 | 510 | 4.545656 | 1533.641 | 0.971796 |
| 7AS6 | 52764 | 21585082 | 409.0873 | 317 | 514 | 3.762411 | 1133.093 | 0.977592 |
| 7AS7 | 51190 | 21098301 | 412.1567 | 274 | 514 | 4.362011 | 1625.58 | 0.970029 |
| 8AS1 | 48849 | 20619379 | 422.1044 | 262 | 510 | 5.573654 | 2034.955 | 0.9619 |
| 8AS2 | 48770 | 20384007 | 417.962 | 203 | 439 | 5.258535 | 2021.609 | 0.962324 |
| 8AS3 | 44098 | 18399510 | 417.2414 | 225 | 503 | 5.292985 | 1886.831 | 0.964021 |
| 8AS4 | 51880 | 21642820 | 417.1708 | 257 | 510 | 5.27387 | 2031.529 | 0.962607 |
| 8AS5 | 51889 | 21488066 | 414.116 | 284 | 493 | 5.708887 | 2067.797 | 0.961688 |
| 8AS6 | 50696 | 21056860 | 415.3555 | 235 | 450 | 6.054447 | 2447.516 | 0.955043 |
| 8AS7 | 36095 | 15018580 | 416.0848 | 373 | 450 | 6.201709 | 2460.788 | 0.9537 |
| 9AS1 | 47215 | 19918801 | 421.8744 | 235 | 510 | 4.501977 | 1731.39 | 0.965293 |
| 9AS2 | 41109 | 17352199 | 422.1022 | 281 | 477 | 5.737845 | 2334.881 | 0.955185 |

| | | | | | | | | |
|---|---|---|---|---|---|---|---|---|
| 9AS3 | 42420 | 17894931 | 421.8513 | 203 | 489 | 5.166586 | 1637.562 | 0.968474 |
| 9AS4 | 39546 | 16658109 | 421.2337 | 202 | 511 | 5.017749 | 1781.54 | 0.966848 |
| 1BS1 | 44466 | 18870353 | 424.3771 | 269 | 455 | 4.291515 | 2047.109 | 0.962183 |
| 1BS2 | 36617 | 15085688 | 411.9859 | 375 | 452 | 5.100266 | 1574.323 | 0.970453 |
| 1BS3 | 39989 | 16492744 | 412.432 | 359 | 452 | 5.406192 | 1607.861 | 0.970877 |
| 1BS4 | 44841 | 18472592 | 411.9576 | 296 | 452 | 5.165863 | 1647.673 | 0.968969 |
| 1BS5 | 49163 | 20609125 | 419.1999 | 283 | 475 | 5.390506 | 2352.908 | 0.953983 |
| 1BS6 | 47355 | 19750283 | 417.0686 | 232 | 466 | 5.767702 | 2707.091 | 0.946915 |
| CX1 | 72469 | 30328435 | 418.5022 | 251 | 460 | 6.20883 | 2626.62 | 0.950802 |
| CX2 | 62098 | 26071403 | 419.8429 | 245 | 522 | 6.097475 | 2363.191 | 0.954407 |
| CZ1 | 51927 | 21665187 | 417.2239 | 229 | 441 | 4.514275 | 2083.609 | 0.95879 |
| CZ2 | 58947 | 24852607 | 421.6094 | 223 | 521 | 6.008745 | 2426.521 | 0.954619 |
| DY1 | 59461 | 24899326 | 418.7505 | 232 | 454 | 4.683268 | 1738.082 | 0.965858 |
| DY2 | 57842 | 24305306 | 420.2017 | 303 | 439 | 6.148294 | 2671.67 | 0.948752 |
| XT1 | 54598 | 22672803 | 415.268 | 231 | 456 | 4.415785 | 2126.311 | 0.957447 |
| XT2 | 60383 | 25373391 | 420.2075 | 245 | 453 | 5.840194 | 1938.742 | 0.96501 |
| 1GP2 | 38396 | 16202544 | 421.9852 | 373 | 456 | 4.381582 | 1723.762 | 0.965858 |
| 1GP3 | 36180 | 15225662 | 420.8309 | 254 | 466 | 5.140201 | 2371.284 | 0.95264 |
| 7GP1 | 49924 | 20612947 | 412.8865 | 317 | 507 | 4.423397 | 1723.006 | 0.965293 |
| 7GP2 | 51271 | 21624918 | 421.7768 | 303 | 432 | 4.19637 | 1242.282 | 0.974553 |
| 7GP3 | 48639 | 20294712 | 417.2518 | 237 | 470 | 4.554037 | 1800.439 | 0.966353 |
| 7GP4 | 58062 | 24203495 | 416.856 | 366 | 437 | 3.891632 | 1500.684 | 0.970877 |
| CJ1 | 54325 | 22838483 | 420.4047 | 298 | 444 | 4.035931 | 650.3958 | 0.988832 |
| CJ2 | 46707 | 19569196 | 418.9778 | 314 | 511 | 3.935175 | 654.2727 | 0.98869 |
| CJ3 | 74098 | 31068462 | 419.2888 | 347 | 444 | 4.077918 | 748.6222 | 0.987771 |

| | | | | | | | | |
|---|---|---|---|---|---|---|---|---|
| CJ4 | 44927 | 18941207 | 421.5996 | 303 | 458 | 4.275643 | 1235.248 | 0.976532 |
| CJ5 | 39335 | 16399481 | 416.9183 | 360 | 432 | 4.850671 | 1189.105 | 0.978794 |
| CJ6 | 52232 | 21708546 | 415.6177 | 302 | 448 | 4.817447 | 1178.484 | 0.979572 |
| CJ7 | 53041 | 22414460 | 422.5874 | 275 | 450 | 4.286674 | 1053.03 | 0.980703 |
| CJ8 | 58750 | 24831085 | 422.6568 | 271 | 448 | 4.994101 | 1256.5 | 0.978299 |
| CJ9 | 43837 | 18505545 | 422.1444 | 230 | 445 | 4.414915 | 638.8182 | 0.989538 |
| CJ10 | 48225 | 20154373 | 417.9238 | 223 | 435 | 4.29138 | 483.1 | 0.992295 |
| CJ11 | 48454 | 20387513 | 420.7602 | 307 | 504 | 4.417303 | 579.3585 | 0.990952 |
| CJ12 | 53981 | 22780752 | 422.0143 | 283 | 468 | 4.654004 | 714.9153 | 0.988973 |
| 7SW1 | 72617 | 30448151 | 419.2978 | 245 | 483 | 4.32451 | 1391.524 | 0.973493 |
| 7SW2 | 69622 | 28914737 | 415.3103 | 274 | 479 | 4.653304 | 1459.581 | 0.970948 |
| 7SW3 | 58226 | 24063877 | 413.284 | 342 | 511 | 5.328393 | 2328.505 | 0.955538 |
| 7SW4 | 69185 | 28986460 | 418.9703 | 337 | 495 | 4.9195 | 2270.218 | 0.958154 |
| 7SW5 | 65192 | 27023412 | 414.5204 | 236 | 518 | 3.276778 | 1227 | 0.978158 |
| 8SW1 | 63700 | 26759135 | 420.0806 | 249 | 479 | 3.704281 | 1347.727 | 0.974482 |
| 8SW2 | 58748 | 24576317 | 418.3345 | 258 | 488 | 4.241372 | 1329.819 | 0.974129 |
| 8SW3 | 66524 | 27456478 | 412.7304 | 232 | 507 | 5.131554 | 2149.531 | 0.958578 |
| 8SW4 | 56592 | 23591208 | 416.8647 | 317 | 468 | 5.371467 | 2681.851 | 0.950025 |
| 8SW5 | 59987 | 25006350 | 416.8628 | 250 | 488 | 5.305262 | 2211.693 | 0.956599 |

**Table S5** Results of the tertiary plot.

| Class | Genus | RW | SW | WW |
|---|---|---|---|---|
| Gammaproteobacteria | Pseudomonas | 41.19% | 2.84% | 9.75% |
| Bacteroidia | Flavobacterium | 3.90% | 6.90% | 7.32% |
| Gammaproteobacteria | Limnohabitans | 7.29% | 5.24% | 4.21% |
| Gammaproteobacteria | Burkholderiaceae_unclassified | 3.29% | 5.62% | 5.94% |
| Gammaproteobacteria | Methylotenera | 10.11% | 1.20% | 3.30% |
| Gammaproteobacteria | Acinetobacter | 3.85% | 1.55% | 5.44% |
| Saccharimonadia | Saccharimonadales_norank | 2.45% | 0.37% | 5.88% |
| Verrucomicrobiae | Prosthecobacter | 1.68% | 1.73% | 3.16% |
| Gammaproteobacteria | Aquabacterium | 0.39% | 2.84% | 3.00% |
| Alphaproteobacteria | Sphingomonadaceae_unclassified | 0.80% | 3.43% | 1.47% |
| Gammaproteobacteria | Acidovorax | 0.48% | 4.35% | 0.71% |
| Verrucomicrobiae | Brevifollis | 1.41% | 2.85% | 0.64% |
| Bacteroidia | Emticicia | 0.64% | 2.33% | 0.65% |
| Bacteroidia | Pedobacter | 0.60% | 1.13% | 1.86% |
| Gammaproteobacteria | Dechloromonas | 0.31% | 1.69% | 1.51% |
| Bacteroidia | Pseudarcicella | 0.54% | 1.88% | 1.00% |
| Gammaproteobacteria | Hydrogenophaga | 1.10% | 1.82% | 0.42% |
| Alphaproteobacteria | Novosphingobium | 0.54% | 1.79% | 0.89% |
| Alphaproteobacteria | Pseudorhodobacter | 0.44% | 1.98% | 0.78% |
| Bacteroidia | Sediminibacterium | 0.63% | 1.09% | 1.44% |
| Gammaproteobacteria | Limnobacter | 0.17% | 1.64% | 1.35% |
| Bacteroidia | NS11-12_marine_group_norank | 0.86% | 1.33% | 0.95% |
| Alphaproteobacteria | Acetobacteraceae_unclassified | 0.07% | 2.95% | 0.01% |
| Gammaproteobacteria | Rheinheimera | 0.35% | 0.77% | 1.87% |
| Parcubacteria | Candidatus_Kaiserbacteria_norank | 0 | 0 | 2.86% |
| Deltaproteobacteria | Peredibacter | 0.45% | 0.76% | 1.60% |
| Gammaproteobacteria | Ideonella | 0.17% | 2.23% | 0.31% |
| Gammaproteobacteria | Methylophilaceae_unclassified | 1.27% | 0.31% | 1.12% |
| Actinobacteria | Micrococcaceae_unclassified | 0 | 2.44% | 0.01% |
| Gammaproteobacteria | GKS98_freshwater_group | 0.29% | 0.38% | 1.74% |
| Alphaproteobacteria | Rhodobacteraceae_unclassified | 0.73% | 1.32% | 0.35% |
| Gammaproteobacteria | Undibacterium | 0.33% | 0.28% | 1.70% |
| Alphaproteobacteria | Caulobacteraceae_unclassified | 0.06% | 1.88% | 0.27% |
| Gammaproteobacteria | Aeromonas | 0.01% | 1.24% | 0.86% |
| Bacteroidia | Fluviicola | 0.64% | 0.96% | 0.39% |
| Verrucomicrobiae | Verrucomicrobiae_norank | 0.13% | 1.72% | 0.00% |
| Campylobacteria | Arcobacter | 0 | 0 | 1.74% |
| Bacteroidia | Chitinophagaceae_norank | 0.53% | 0.66% | 0.36% |
| Gammaproteobacteria | Cupriavidus | 0 | 1.36% | 0.10% |
| Verrucomicrobiae | Pedosphaeraceae_norank | 0.14% | 1.16% | 0.10% |

| | | | | |
|---|---|---|---|---|
| Gammaproteobacteria | Polaromonas | 1.23% | 0.14% | 0.00% |
| Acidobacteriia | Paludibaculum | 0.22% | 1.10% | 0.01% |
| Gammaproteobacteria | Malikia | 0.08% | 0.03% | 1.21% |
| Bacteroidia | 37-13_norank | 0.57% | 0.59% | 0.14% |
| Gammaproteobacteria | Massilia | 0.48% | 0.16% | 0.61% |
| Gammaproteobacteria | Lautropia | 0.03% | 1.16% | 0.01% |
| Alphaproteobacteria | Sphingobium | 0.13% | 0.81% | 0.22% |
| WWE3 | WWE3_norank | 0.01% | 0 | 1.13% |
| Alphaproteobacteria | Rhodobacter | 0.11% | 0.82% | 0.19% |
| Gammaproteobacteria | Perlucidibaca | 0.78% | 0.21% | 0.11% |
| Alphaproteobacteria | Phenylobacterium | 0.08% | 0.62% | 0.36% |
| Gammaproteobacteria | Polynucleobacter | 0.23% | 0.16% | 0.66% |
| Deltaproteobacteria | 053A03-B-DI-P58_norank | 1.01% | 0 | 0 |
| Alphaproteobacteria | Roseomonas | 0.06% | 0.88% | 0.07% |
| Alphaproteobacteria | Alphaproteobacteria_norank | 0.01% | 0.94% | 0.01% |
| Bacteroidia | Runella | 0.07% | 0.77% | 0.06% |
| Parcubacteria | Parcubacteria_unclassified | 0 | 0 | 0.88% |
| Oxyphotobacteria | Chloroplast_norank | 0 | 0.05% | 0.83% |
| Deltaproteobacteria | Bacteriovorax | 0.10% | 0.15% | 0.57% |
| Gammaproteobacteria | Ramlibacter | 0.01% | 0.69% | 0.04% |
| Ignavibacteria | OPB56_norank | 0.57% | 0.03% | 0.13% |
| Deltaproteobacteria | OM27_clade | 0 | 0.71% | 0.01% |
| Parcubacteria | Candidatus_Moranbacteria_norank | 0 | 0 | 0.67% |
| Bacteroidia | Crocinitomix | 0.55% | 0 | 0 |
| | others | 5.79% | 13.96% | 15.00% |

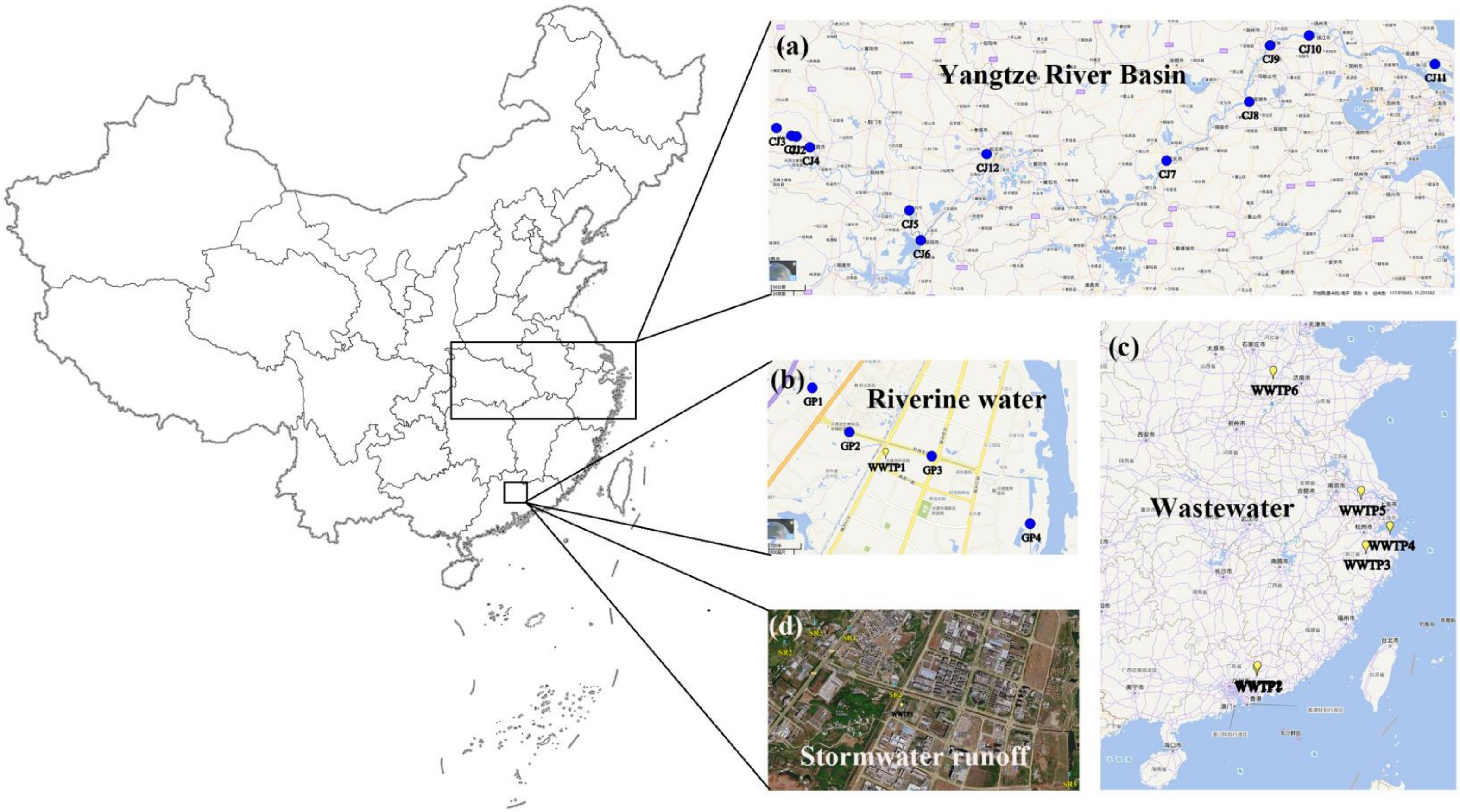


**Fig. S1** Sampling sites of the riverine water from the Yangtze River Basin (a) and Gaopu Stream (b), wastewater from the six full scale wastewater treatment plants (WWTPs, c) and stormwater runoff (d).

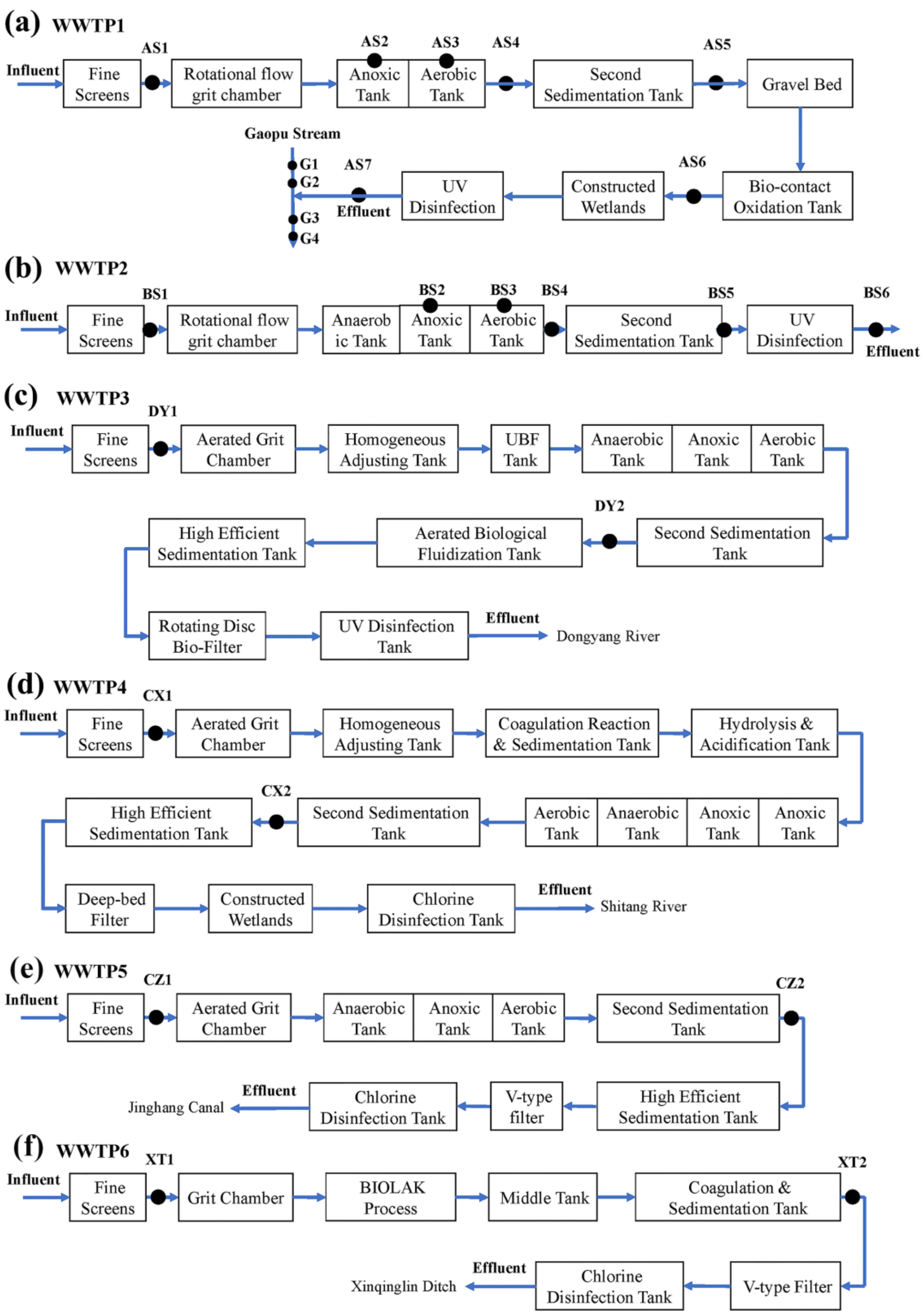


**Fig. S2** The treatment processes of the six full-scale WWTPs.

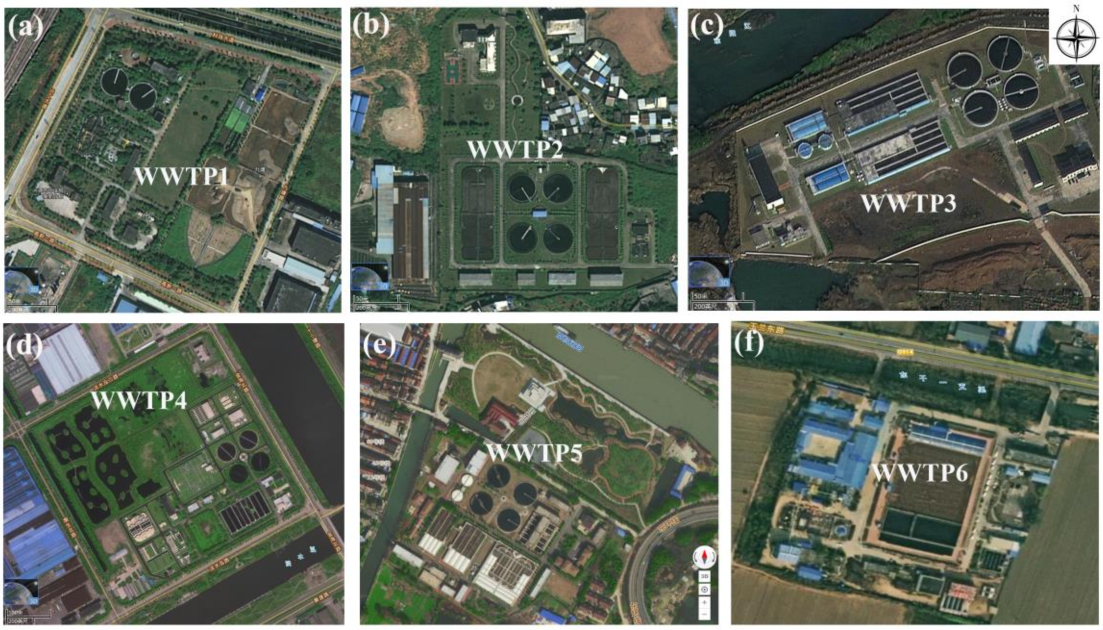


**Fig. S3** The aerial views of the six WWTPs.

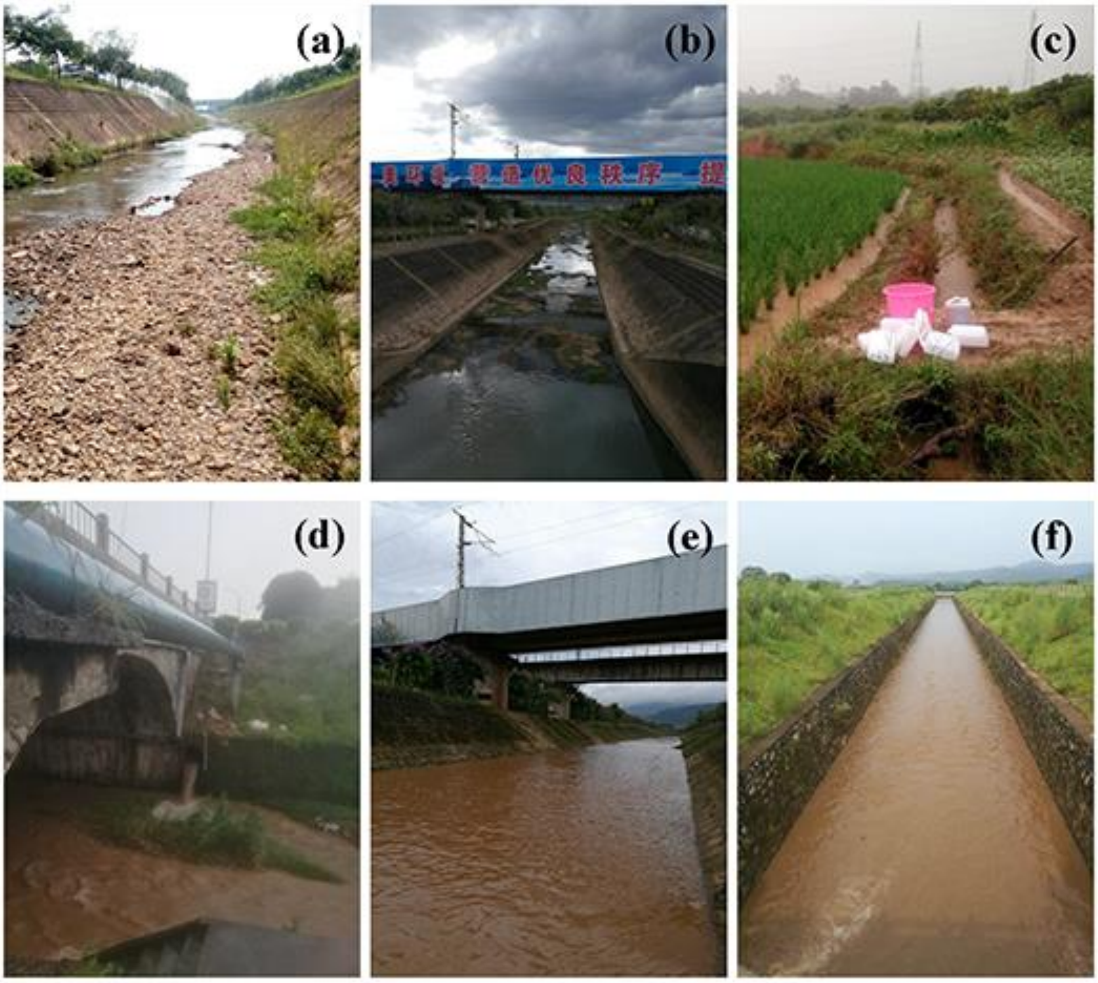


**Fig. S4** The pictures show the sampling sites of stormwater runoff. In the normal season, the average flow of the stream is less than 0.8 $m^3/s$ (a and b). The agricultural zone (c), the rain water pipe from the residential area (d), the sampling site SR4 (e) and site SR5 (f) in the downstream are presented.

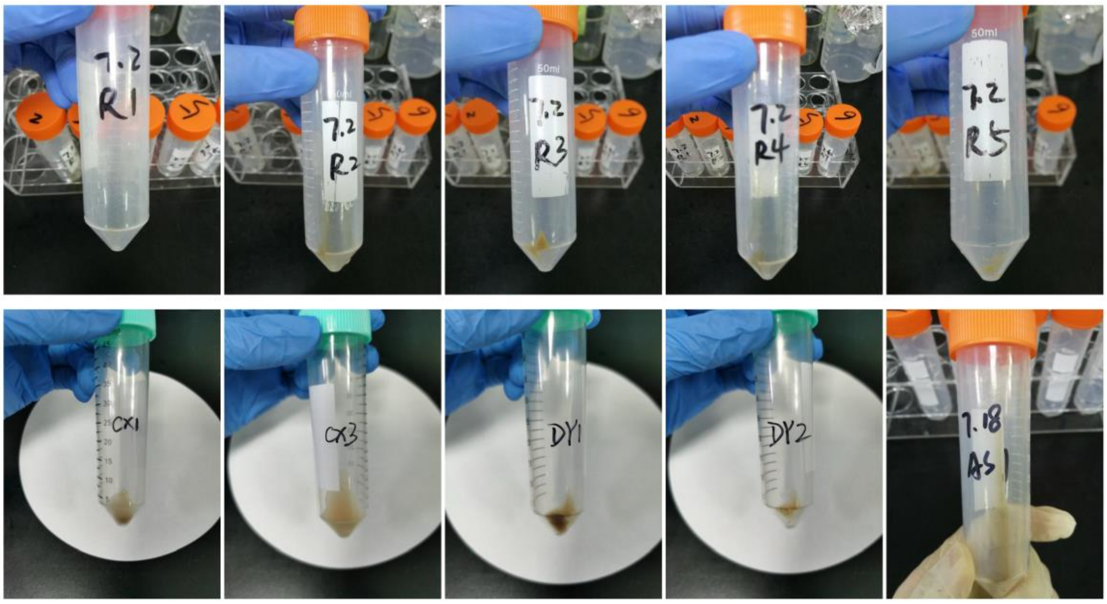


**Fig. S5** The obtained extracellular precipitates after centrifuge from stormwater runoff and wastewater samples.

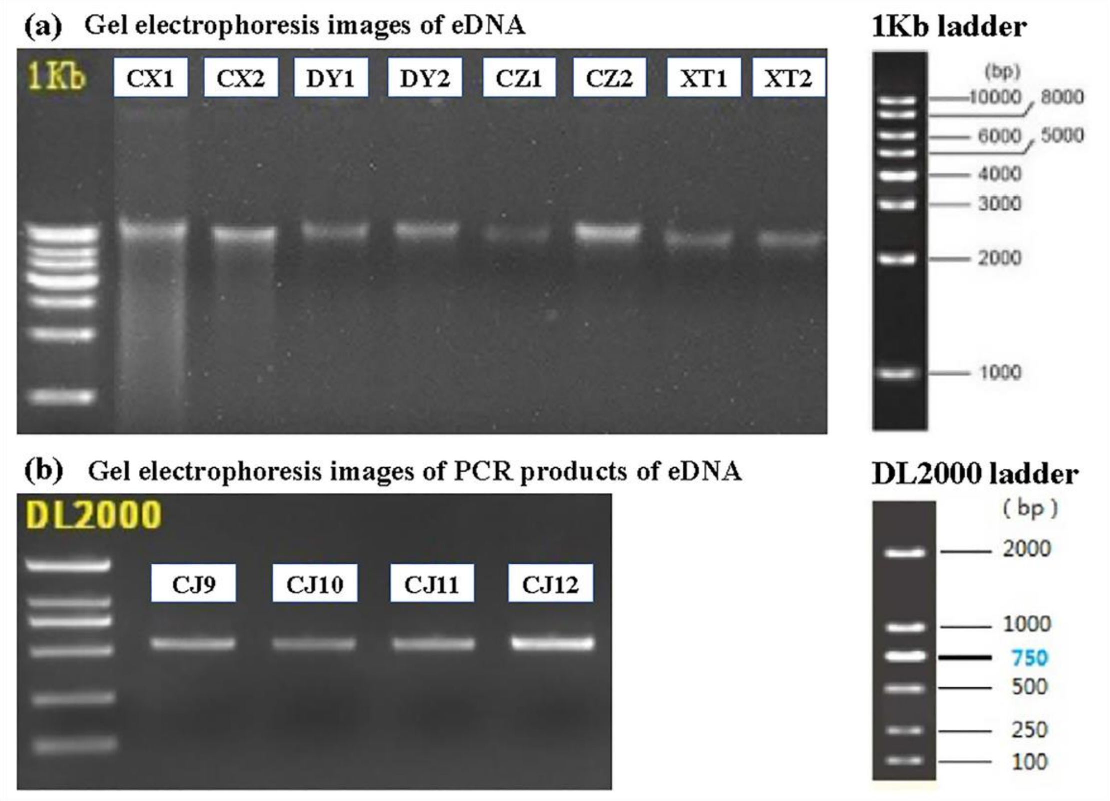


**Fig.** S6 Gel electrophoresis images of eDNA (a) and corresponding PCR products (b).

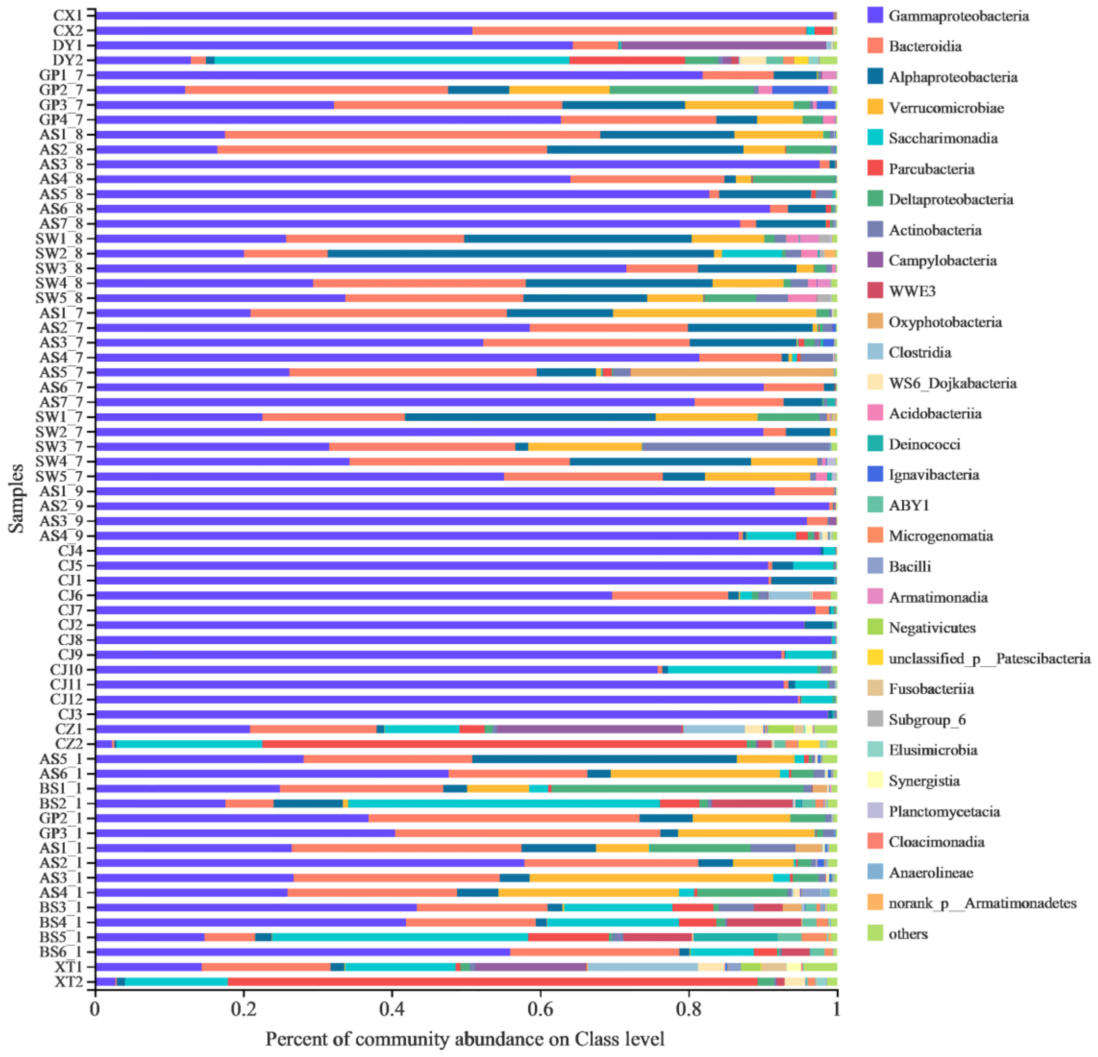


**Fig. S7.** The relative abundances of microbial communities of all eDNA samples on Class level. Those relative abundances less than 1% were combined into Others.

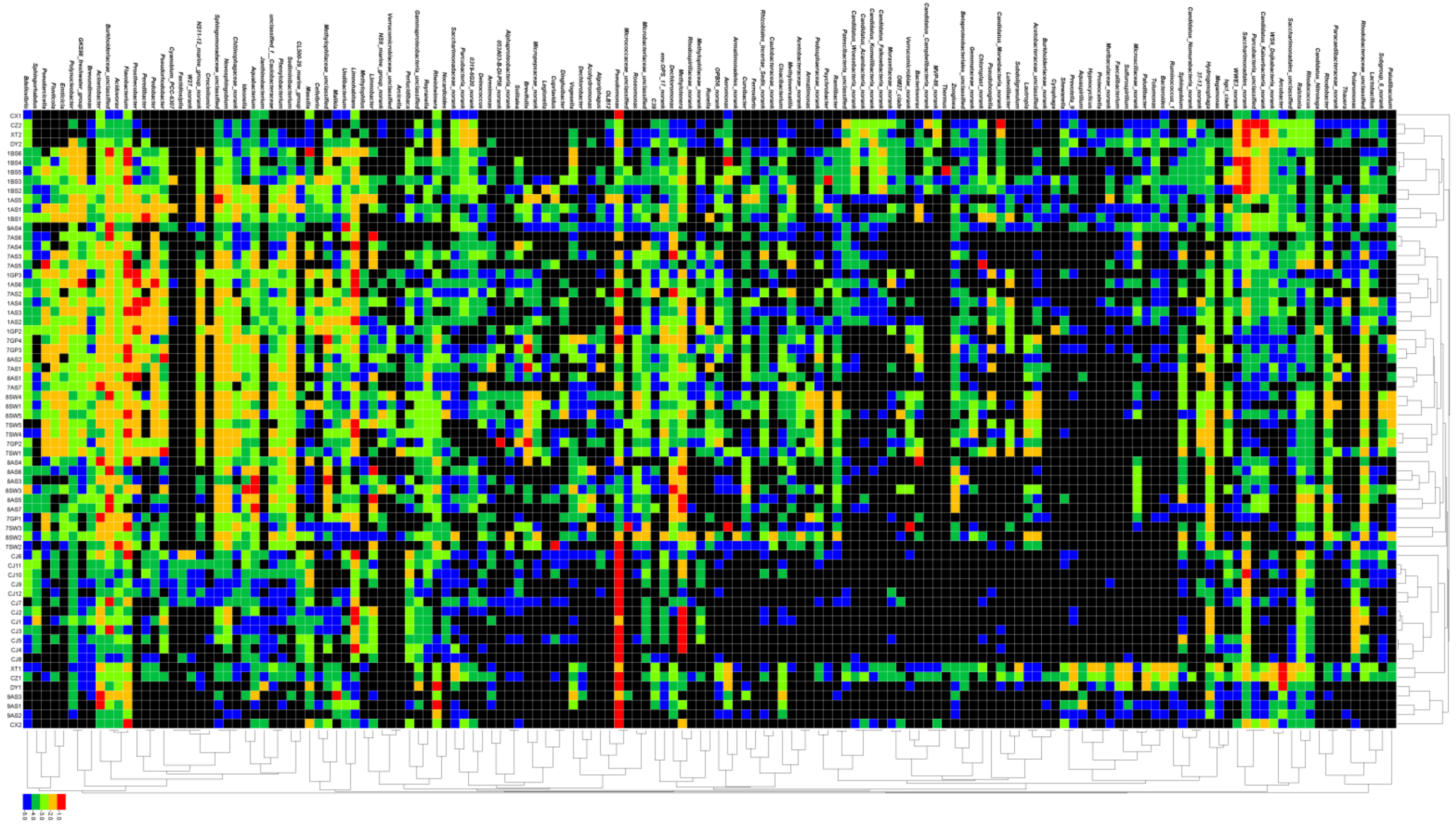

**Fig. S8.** The heatmap shows the relative abundances (lg transformed) of the abundant genera of the eDNA samples (> 1.0%). Black means not detected.

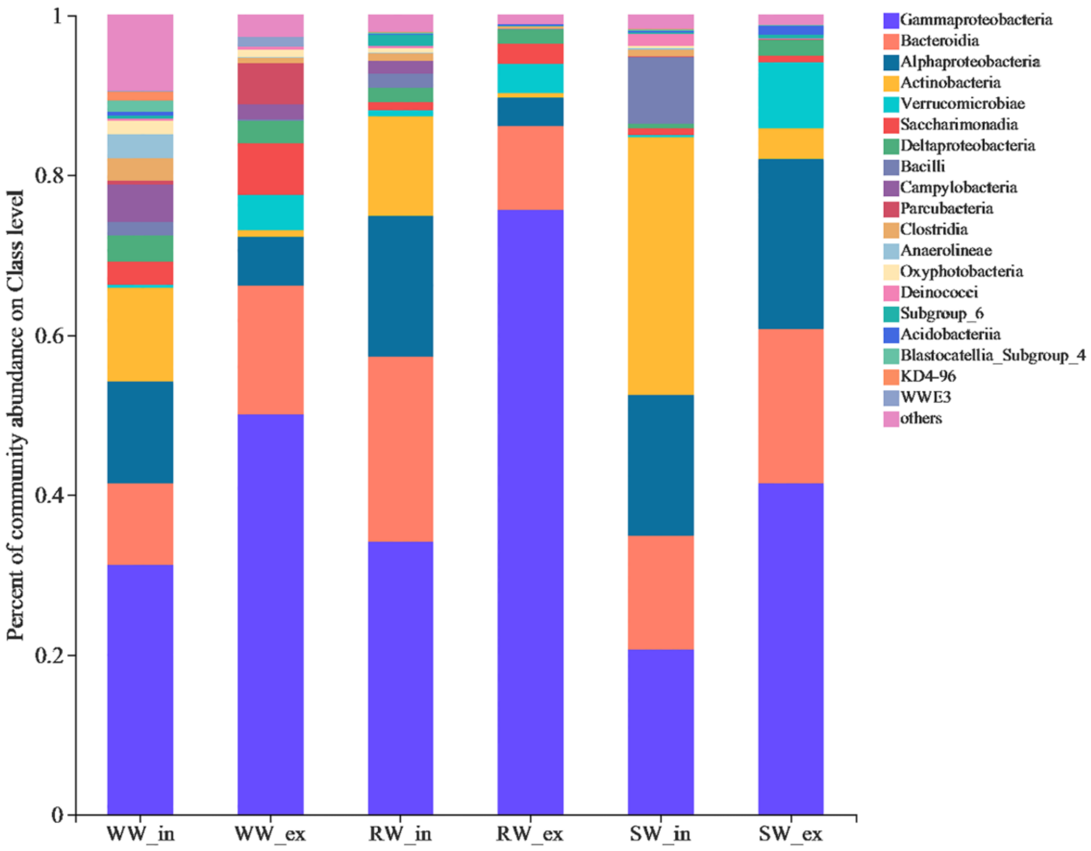


**Fig. S9.** The relative abundances of microbial communities of the iDNA and eDNA samples on Class level.

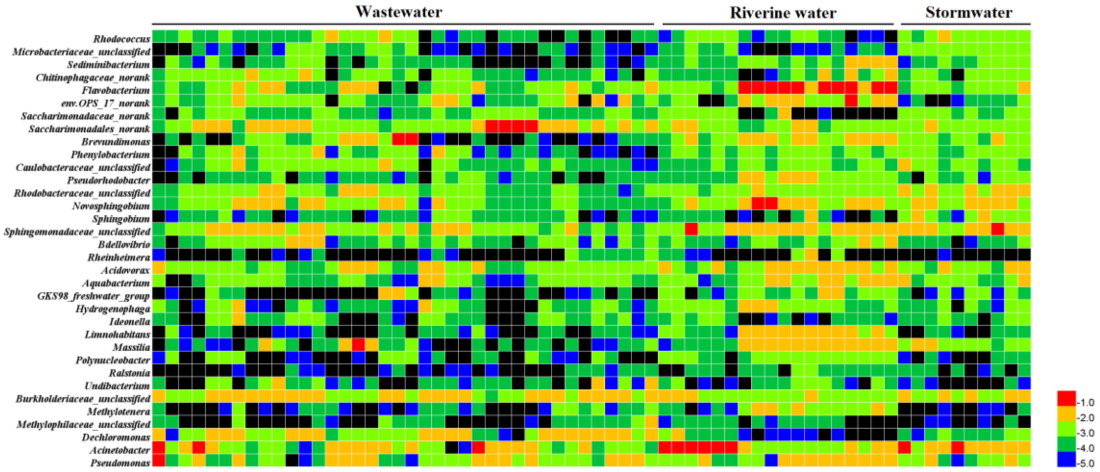


**Fig. 10** The heatmaps show the relative abundances (lg transformed) of bacteria (genus level) from the iDNA samples.

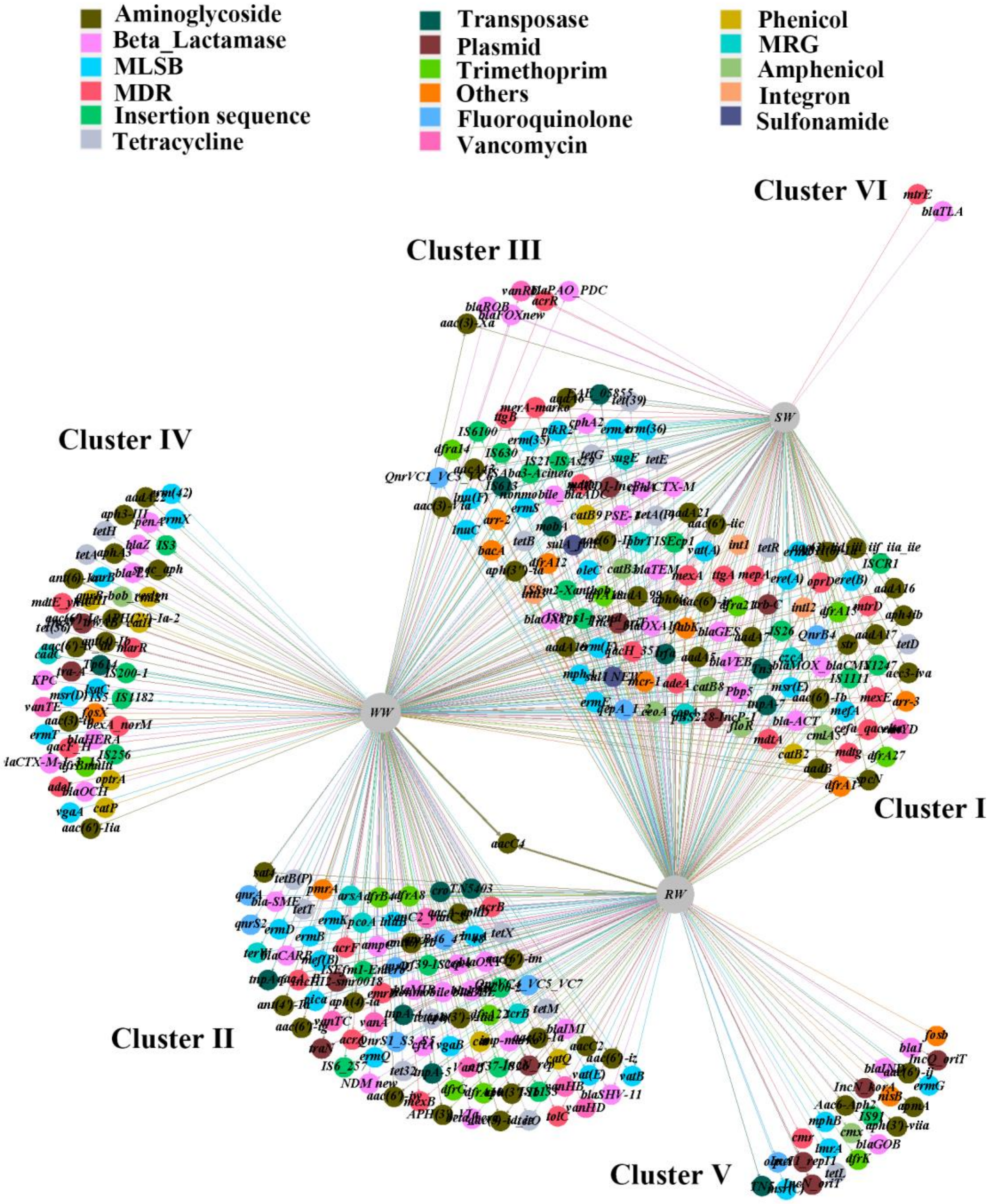


**Fig. S11** Bipartite network analysis presents the exact shared and unique resistance genes among the three water compartments.

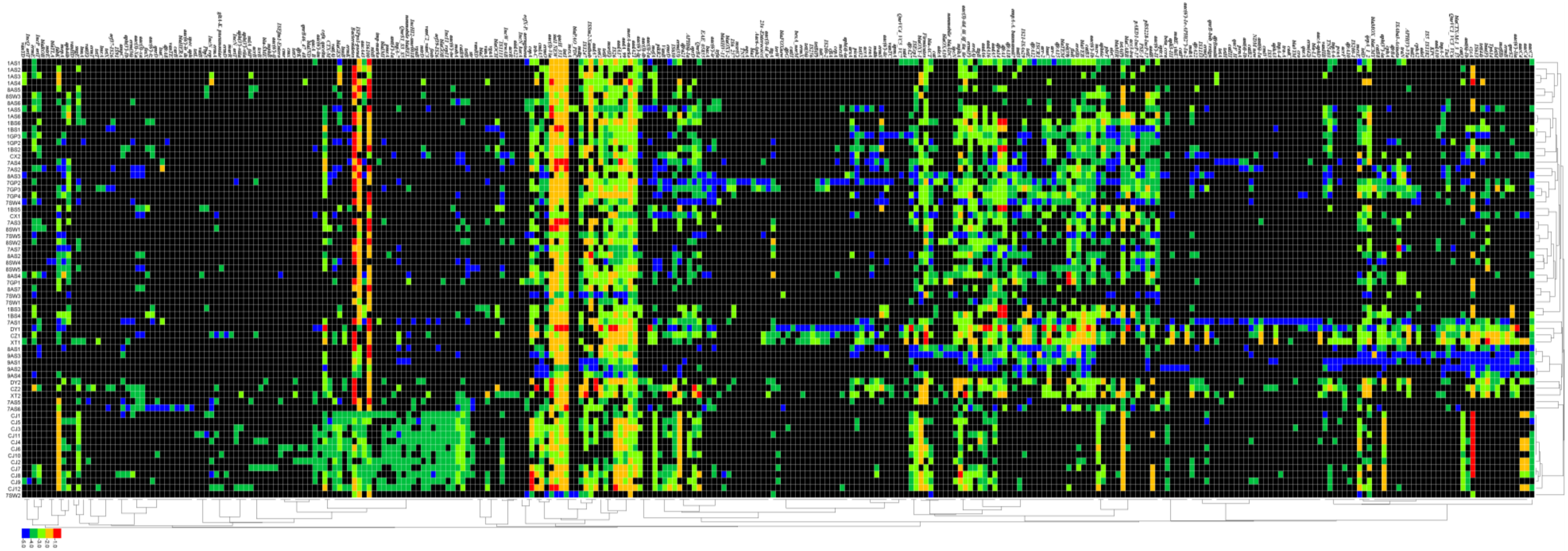

**Fig. S12** The heatmap shows the relative abundances (lg transformed) of all detected extracellular resistance genes. Black means not detected.
